# Load-dependent RGD-context sensing *via* αV-class integrin reprograms cellular adhesion and mechanics within seconds

**DOI:** 10.64898/2025.12.09.693059

**Authors:** Upnishad Sharma, Jakob Reber, Jonne Helenius, Cara Buchholz, Reinhard Faessler, Nico Strohmeyer, Daniel J. Müller

## Abstract

The cellular ability to biophysically and biochemically recognize extracellular matrix proteins is fundamental to adhesion, mechanics, migration, and morphogenesis, and influences homeostasis and disease. However, the mechanisms underlying integrin-mediated mechanosensing of the arginine-glycine-aspartic acid (RGD)-motif of vitronectin and fibronectin remain elusive. Here, we discover that within seconds of sensing vitronectin, αV-class integrins initiate and strengthen adhesion biphasically through complementary mechanotransduction pathways, which rely on the catch bond behavior of single αVβ3 integrins. The first adhesion phase requires αVβ3 and αVβ5 integrin-associated actomyosin and FAK activity, while αVβ5 integrin additionally requires clathrin-mediated endocytosis. With elevating mechanical load, the second phase requires αVβ3 integrin-directed Arp2/3, cSrc, and PI3K signaling that dominates αVβ5 integrin in organizing the consensus adhesome on vitronectin. Simultaneously, αVβ5 integrin regulates the mechanical stiffening of fibroblasts. Thus, αV-class integrins exhibit rapid RGD-motif- and β-subunit-specific programs to synergistically guide mammalian cell adhesion and mechanics upon encountering diverse extracellular environments.

## Main

Mammalian cells adhere to the extracellular matrix (ECM) using highly dynamic and hierarchical integrin-based adhesion complexes that establish an ECM-receptor-adaptor-cytoskeleton axis^1-4^. Integrins are obligatory heterodimeric transmembrane receptors that enable adherent cells to sense and interpret various biochemical, structural and mechanical cues in their immediate environment^5, 6^. Different classes of integrins sense diverse ECM proteins like fibronectin or vitronectin, and recruit and/or activate proteins to assemble adhesion complexes^7-10^. Juxtaposition of these varied cues is critical for biological processes such as wound healing and development^11-14^. αV-class (αVβ1, αVβ3, αVβ5, αVβ6, and αVβ8) integrins bind to the arginine-glycine-aspartic acid (RGD)-motif of ECM proteins and govern early morphogenesis^15^, osteogenesis, lipid biogenesis^16^ besides being involved in several disease conditions including fibrosis^17^, tumor metastasis, and age-related macular degeneration^10, 18-24^.

Upon ligand binding, integrins assemble nascent adhesions that mature into more complex adhesion complexes as the actomyosin-mediated contractile forces continuously increase, on timescales of minutes to hours^4, 25-27^. Additionally, intra- or extracellular tensile forces influence the assembly and dynamics of adhesion complexes^1, 11, 14, 25, 28-33^. How rapidly αV-class integrins recognize the RGD-motif of the key ECM proteins, fibronectin and vitronectin, and respond at the onset of cell adhesion has not been addressed, mainly due to the limited temporal and force resolution of the biophysical techniques applied^1, 34-36^. Moreover, it is debated whether cell adhesion to vitronectin is mechanosensitive, while integrin mechanotransduction triggered by fibronectin is well studied^27, 37-41^. How αV-class integrins may relay this contextualization of the RGD-motif under varying mechanical load, such as experienced within tissues, demands enquiry. Although, the effects of cell-generated forces on different αV-class and β1-class integrins has been studied from few minutes to several hours^34, 42^, the effect of dynamic force-loading during early cell adhesion (<120 s) to vitronectin remains elusive. Thus, exploring the even earlier time ranges of adhesion is fundamental to understanding how cells employ integrins to mechanosense the ECM and initiate cellular responses.

Here, we explore the β-subunit dependence of αV-class integrins in initiating and mechanically strengthening cell adhesion to vitronectin and fibronectin. Employing an interdisciplinary approach including atomic force microscopy (AFM)-based single-cell force spectroscopy (SCFS), mass spectrometry-based adhesion proteomics, chemical perturbations, and fluorescence microscopy, we characterize the early and late adhesion of genetically engineered mouse embryonic fibroblasts. We find that αV-class integrins distinguish the different RGD-context in fibronectin and vitronectin within seconds and that this contextualization leads them to strengthen individual integrin-bonds as well as cell adhesion either monophasically or biphasically in response to mechanical load. Thereby, αV-class integrins facilitate the unique mechanosensitive biphasic adhesion response to vitronectin through two different integrin β-subunits, each of which using specific components of the actomyosin cytoskeleton and biochemical signaling pathways to establish substrate-specific cell adhesion and mechanical responses.

## Results

### αV-class integrins distinguish context of RGD-motif under mechanical load

To understand whether αV-class integrins distinguish and hence respond to the RGD-motif context, we used mouse embryonic kidney fibroblasts (pKO-αV, pan integrin knockout cells with β2^-/-^, β7^-/-^, β1^-/-^, αV^-/-^, αV reintroduced) expressing two αV-class integrins, αVβ3 and αVβ5 at similar levels on the cell surface^27^. Mass spectrometry analysis of the adhesion assemblies (adhesome) of these fibroblasts cultured on vitronectin- or fibronectin-substrates for 45 min showed substrate-dependent differences (Fig. 1a). On fibronectin, pKO-αV fibroblasts enriched different collagens (Col1a1, Col1a2, Col3a1, Col12a1, Col18a1, etc.), whereas pKO-αV fibroblasts cultured on vitronectin enriched several components of the complement system (C3, C4b, C8b, C8g, Cfb, etc.). This considerable difference suggests that within 45 min of adhesion, pKO-αV fibroblasts differentially respond to the RGD-motif context of vitronectin and fibronectin.

**Fig. 1:**
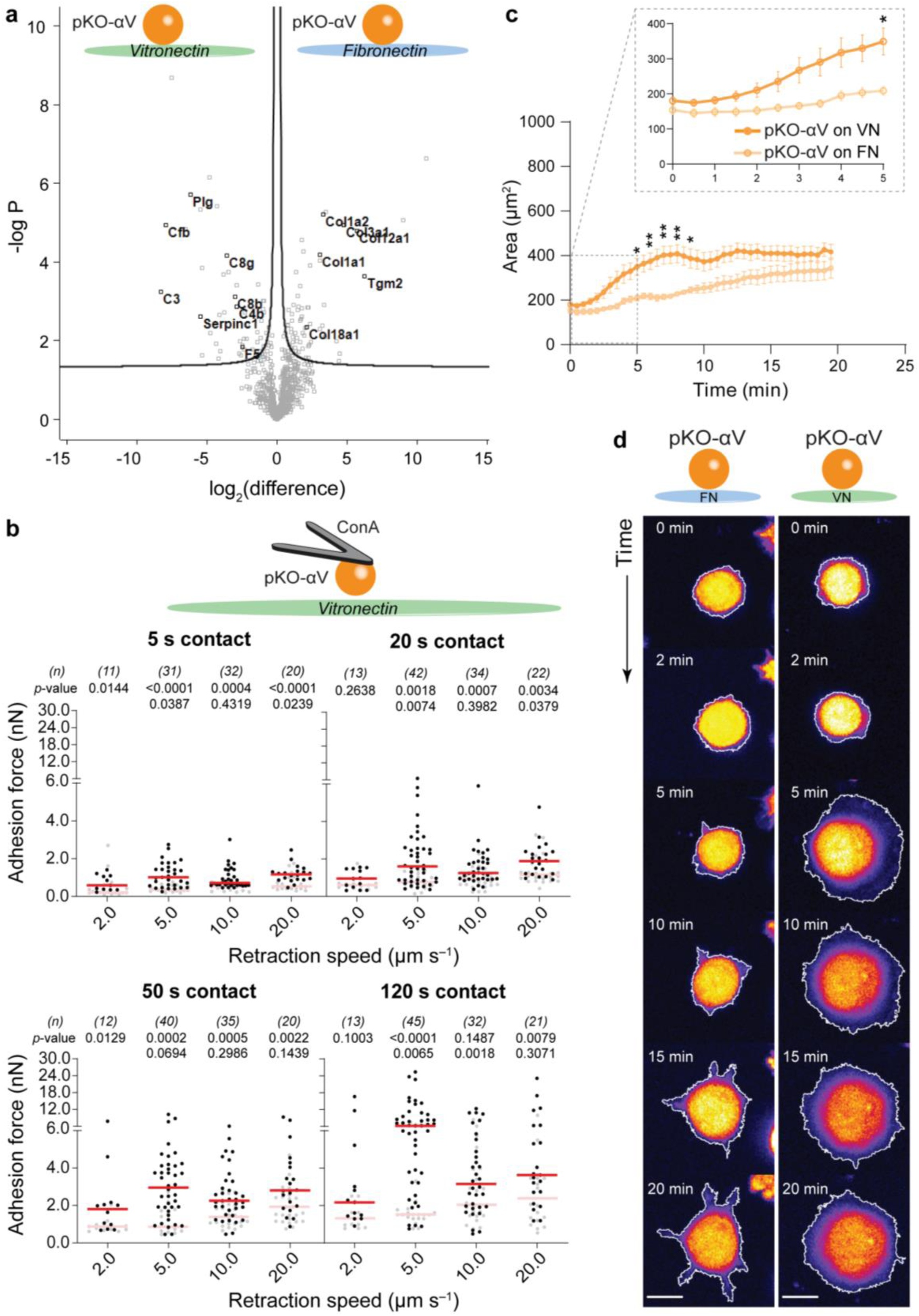
αV-class integrins distinguish RGD context under mechanical load. **a**, Composition of adhesomes isolated from pKO-αV fibroblasts adhering to fibronectin- or vitronectin-coated glass substrates for 45 min. Volcano plot of mass spectrometry (MS) analysis show the abundance of ECM-related proteins. Black lines represent the boundaries used to identify protein candidates corresponding to a false-discovery rate of 0.1 and a penalty factor for sample-wide standard deviation of 0.1. Data is representative of four independent replicates. **b**, Adhesion force of pKO-αV fibroblasts to vitronectin-coated glass increases ‘biphasically’ with retraction speed and contact time. The biphasic increase in adhesion force becomes most prominent at 120 s contact time. Each dot represents the measurement of a single fibroblast, red bars correspond the median. *p*-values were obtained using two-tailed Mann-Whitney U-test. *p*-values in the top row compare cell adhesion force on vitronectin (black) to fibronectin (lightened). *p*-values in the bottom row compare data for a given retraction speed to that of the preceding retraction speed (e.g., 5 µm s^-1^ with 2 µm s^-1^). *n* denotes the number of fibroblasts characterized in three different experimental days. **b**, Spreading kinetics of pKO-αV fibroblasts on fibronectin (FN) or vitronectin following their first contact to the substrate. Confocal microscopy images analyzed were acquired every 30 s. Inset shows the spreading dynamics of fibroblasts on vitronectin and fibronectin within the first 5 min after attachment in better detail. Each data point presents the mean ± SE from > 10 fibroblasts. * *p*-value < 0.05; ** *p*-value < 0.01 obtained using two-tailed Mann-Whitney U-test. **d**, Time-stamped confocal microscopy of pKO-αV fibroblasts stained with CellTracker™ dye (shown in pseudocolor) and spreading on fibronectin or vitronectin. White outlines represent the masks used for calculating the projected cell areas used in (**c**). Scale bars, 10 µm.

Next, we characterized whether this contextualization of the RGD-motif occurs while fibroblasts establish adhesion and are exposed to mechanical load. Utilizing AFM-based SCFS^43-46^ (Extended Data Fig. 1a,b), we quantified the adhesion forces of pKO-αV fibroblasts to fibronectin- or vitronectin-coated substrates for contact times between 5 – 120 s, by detaching them at different retraction speeds of 2 to 20 µm s^-1 37, 47^,which simulated mechanical loading rates comparable to those occurring in tissues, such as interstitial fluid flows^48^. In case of vitronectin, we observed a rapid increase in cell adhesion force between 2 and 5 µm s^-1^ (Fig. 1b). At 10 µm s^-1^, however, the adhesion force reduced and thereafter increased more gradually until ≤ 20 µm s^-1^. We term this rapid increase in adhesion force followed by a decrease and thereafter by a gradual increase, ‘biphasic adhesion response’.

This biphasic adhesion response became more prominent with increasing contact time. Further, confirmation of the biphasic cell adhesion response was achieved by a more detailed probing the retraction speed regime of 3, 6 and 8 µm s^-1^ (Extended Data Fig. 2a). Markedly, pKO-αV fibroblasts showed a monophasic adhesion response to fibronectin with increasing retraction speeds (Extended Data Fig. 2b), thereby confirming the specificity of the biphasic adhesion response of fibroblasts to vitronectin.

To characterize the cell morphological effect of the RGD-context, we performed timelapse confocal microscopy of pKO-αV fibroblasts adhering to vitronectin or fibronectin during early stages of cell spreading within 20 min (Fig. 1c,d). On vitronectin, pKO-αV fibroblasts spread over larger areas (∼400 µm^2^) and faster (within 10 min at a rate of 36.45 µm^2^ min^-1^) than on fibronectin (∼350 µm^2^, within 20 min at 2.46 µm^2^ min^-1^). Together the data emphasizes the necessity to better elucidate the early (< 5 min) phases of RGD-contextualization and αV-class integrin dependent cell adhesion.

### Integrin β-subunits dictate fibroblasts biphasic adhesion response

The presence of both αVβ3 and αVβ5 integrins in pKO-αV fibroblasts hinders the attribution of vitronectin-specific biphasic adhesion response to both or one of the two heterodimers. To delineate the contribution from αVβ3 and αVβ5 integrins, we performed a CRISPR/Cas9-mediated deletion of the gene encoding for integrin β3 (*Itgb3*)- or β5 (*Itgb5*)-subunit in pKO-αV fibroblasts, which resulted in pKO-αVβ5 and pKO-αVβ3 fibroblasts, respectively (Fig. 2a). Thereafter, pKO-αVβ3 and pKO-αVβ5 fibroblasts were sorted using flow cytometry to obtain cells with integrin expressions similar to pKO-αV fibroblasts (Fig. 2a). The sorted fibroblasts were then subjected to SCFS as described above, to quantify their early adhesion to vitronectin or fibronectin. To vitronectin, pKO-αVβ3 fibroblasts rapidly increased adhesion force upon increasing the retraction speed from 2 to 5 µm s^-1^, though with lower force compared to pKO-αV fibroblasts (Fig. 2b). The adhesion force considerably reduced at retraction speed of 10 µm s^-1^ and remained low at 20 µm s^-1^. On the contrary, pKO-αVβ5 fibroblasts increased the adhesion force to vitronectin with increasing the retraction speed from 2 to 5 µm s^-1^ and plateaued with high adhesion force at higher retraction speeds (Fig. 2c). This suggests that αVβ3 and αVβ5 integrin differentially contribute to the biphasic cell adhesion response upon mechanical load. Moreover, this biphasic adhesion in response to increasing retraction speed, which is mediated through αVβ3 and αVβ5 integrins, is unique to vitronectin and not observed on fibronectin (Fig. 2d,e and Extended Data Fig. 3).

**Fig. 2:**
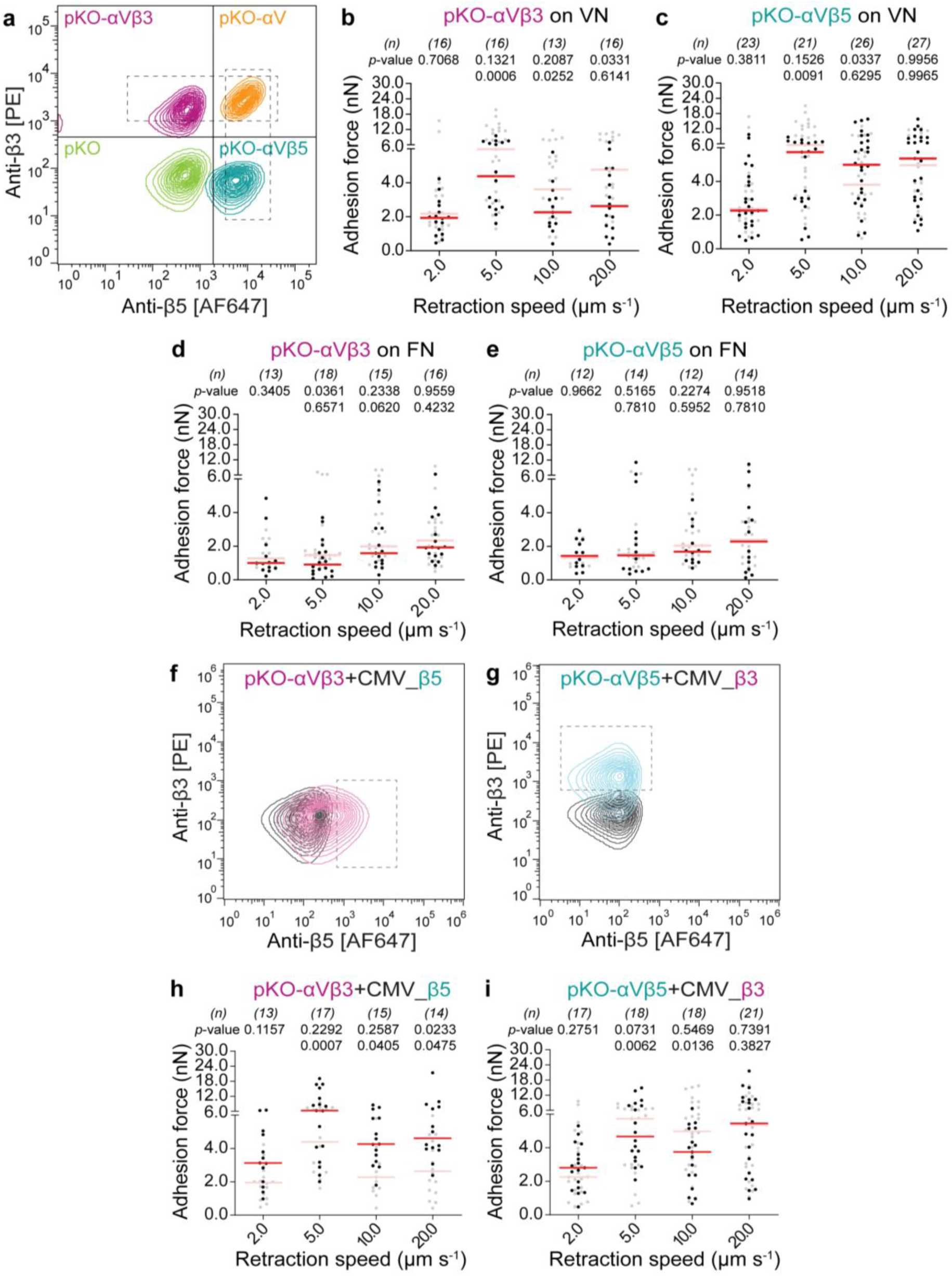
Fibroblasts adhering to vitronectin respond biphasically to mechanical load in an integrin β-subunit specific manner. **a**, Flow cytometry analysis of pKO, pKO-αVβ5, pKO-αV, and pKO-αVβ3 fibroblasts. Fibroblasts were doubly labelled with anti-β3-subunit and anti-β5-subunit antibodies (Methods). (**b**-**e**) Adhesion force of pKO-αVβ3 fibroblasts and pKO-αVβ5 fibroblasts to vitronectin (**b** and **c**) and to fibronectin (**d** and **e**) in dependence of the retraction speed. All fibroblasts adhered to the substrate for 120 s. **b**, The biphasic adhesion response of pKO-αVβ3 fibroblasts to vitronectin is similar to that of pKO-αV fibroblasts (lightened data), but overall shows lower force. **c**, Biphasic adhesion response of pKO-αVβ5 fibroblasts to vitronectin, which is distinct from pKO-αV fibroblasts (lightened data). The adhesion force plateaus at ≥ 5 µm s^-1^. **d**, pKO-αVβ3 fibroblasts adhering to fibronectin show a monophasic adhesion response to the increasing retraction speed, which is similar to pKO-αV fibroblasts (lightened data). The lightened data in (**b**) and (**c**) compare adhesion force of pKO-αV fibroblasts to vitronectin (Extended Data Fig. 3a). The lightened data in (**d**) and (**e**) compare adhesion force of pKO-αV fibroblasts to fibronectin (Extended Data Fig. 3b). **e**, pKO-αVβ5 fibroblasts adhering to fibronectin show a monophasic adhesion response with increasing retraction speed, which is similar to pKO-αV fibroblasts (lightened data). **f**, Flow cytometry sorting scheme to confirm CMV_β5 rescued pKO-αVβ3 fibroblasts (pink) compared to isotype control (grey). The grey box represents the fibroblasts sorted for similar surface integrin expression to pKO-αV fibroblasts and these were used for further experimentation. **g**, Flow cytometry sorting scheme to confirm CMV_β3 rescued pKO-αVβ5 fibroblasts (blue) compared to isotype control (grey). The grey box represents the fibroblasts sorted for similar surface integrin expression to pKO-αV fibroblasts and these were used for further experimentation. **h**, pKO-αVβ3 fibroblasts rescued for integrin β5 expression (dark red bar, black dots) show a distinct biphasic increase of adhesion force with increasing retraction speed. The lightened data are adhesion forces of non-rescued pKO-αVβ3 fibroblasts. **i**, pKO-αVβ5 fibroblasts rescued for integrin β3 expression (dark red bar, black dots) show a restored biphasic increase of adhesion force with increasing retraction speed. The lightened data show adhesion forces of non-rescued pKO-αVβ3 fibroblasts. All the SCFS data was recorded at 120 s contact time on vitronectin substrate. Red bars in **b-e**, **g**, and **i** represent the median. *p*-values were obtained using two-tailed Mann-Whitney U-test. *p*-values in the top row compare data to control (lightened) data. *p*-values in the bottom row compare data for a given retraction speed to the preceding retraction speed (e.g., 5 µm s^-1^ with 2 µm s^-1^). *n* denotes the number of fibroblasts measured.

### Individual integrin β-subunits govern different features of biphasic adhesion response

To explore how integrin β3- and β5-subunits facilitate the biphasic adhesion response of fibroblasts we restored their surface expression in pKO-αVβ5 and pKO-αVβ3 fibroblasts, respectively (Fig. 2f,g). Upon rescuing the respective integrin β-subunit, we found the biphasic adhesion response of pKO-αVβ3 as well as pKO-αVβ5 fibroblasts to be similar to pKO-αV fibroblasts (Fig. 2h,i). Interestingly, rescuing the β5-subunit in pKO-αVβ3 fibroblasts led to a higher cell adhesion force, especially at higher retraction speeds (Fig. 2h). On the other hand, rescuing the β3-subunit in pKO-αVβ5 fibroblasts considerably lowered the cell adhesion force at 10 µm s^-1^ compared to 5 µm s^-1^ (Fig. 2i). Complementary to the genetic manipulation, the inhibition of the integrin β3-subunit in pKO-αV fibroblasts by the αVβ3 integrin antagonist P11 corroborated the specific contribution of the β5-subunit to the biphasic cell adhesion response, though at lower overall adhesion force (Extended Data Fig. 4).

Additionally, the data suggests that the αV-subunit, being the only unaltered subunit among the fibroblasts, dominates the considerable increase in cell adhesion force between 2 µm s^-1^ and 5 µm s^-1^ (Fig. 1b, Extended Data Fig. 2a and Fig. 2h,i). Based on this observation we infer that the rapid increase in cell adhesion force between 2 µm s^-1^ and 5 µm s^-1^ was conserved for pKO-αV, pKO-αVβ3, and pKO-αVβ5 fibroblasts, despite their different integrin β-subunit composition. Because the cell adhesion force between 5 µm s^-1^ and 10 µm s^-1^ plateaued for integrin β5-subunits and decreased for integrin β3-subunits the mechanism critically depends on the β-subunit. Therefore, the repression and rescue of the β-subunit expression in pKO-αV fibroblasts delineated the role of integrin α- and β-subunits in responding to mechanical load.

### β3- and β5-subunits modulate single integrin-vitronectin catch bonds

Next, we tested whether the cell adhesion response to varying mechanical load (retraction speed) arises from the single integrin-vitronectin (receptor-ligand) bond characteristics. Thereto, we analyzed the unbinding events of single integrin-vitronectin bonds detected in SCFS (Methods, Extended Data Fig. 5a)^45, 46, 49^. αVβ5 integrin in pKO-αVβ5 fibroblasts showed the highest binding probability to vitronectin of 0.76 ± 0.03 (mean ± SE), whereas αVβ3 integrin in pKO-αVβ3 fibroblasts showed vitronectin-binding probabilities of 0.68 ± 0.02. pKO-αV fibroblasts, showed lowest vitronectin-binding probability of 0.59 ± 0.03. Thereafter, we analyzed the forces and lengths (lifetimes) of tethers pulled from the cell membrane (Extended Data Fig. 5b), which were recorded upon mechanically detaching fibroblasts from vitronectin by SCFS and describe the biophysical properties of single integrin-vitronectin bonds^50^. The median tether force of pKO-αV, pKO-αVβ3, and pKO-αVβ5 fibroblasts increased with the retraction speed (Extended Data Fig. 5c-e). However, only pKO-αVβ5 fibroblasts rapidly increased the tether force by >60 pN between 5 µm s^-1^ (31.36 ± 1.98 pN; median ± 95% CI) and 10 µm s^-1^ (98.89 ± 9.35 pN) retraction speed, indicating altered cell membrane properties^49^. The pKO-αV and pKO-αVβ3 fibroblasts showed a similar trend and increased the tether force with retraction speed, though pKO-αV fibroblasts showed higher tether forces (by >10 pN) compared to pKO-αVβ3 fibroblasts except for 20 µm s^-1^ retraction speed (Extended Data Fig. 5c,d). The median tether lifetime at 2, 5, and 10 µm s^-1^ retraction speed for the pKO-αVβ5 fibroblasts were higher compared to the pKO-αVβ3 fibroblasts (Extended Data Fig. 5g,h).

Lastly, we analyzed the rupture force of single integrin-vitronectin bonds, which had also been recorded by SCFS upon detaching fibroblasts from vitronectin (Extended Data Fig. 5b). The median rupture force of integrin-vitronectin bonds for pKO-αVβ5 fibroblasts for all retraction speeds was higher than pKO-αV fibroblasts (Extended Data Fig. 5i,k,l,n), while pKO-αVβ3 fibroblasts displayed the lowest rupture forces at all retraction speeds (Extended Data Fig. 5j,m). The rupture force of single αVβ3 integrins increased with the mechanical load non-linearly (Extended Data Fig. 5m). This indication of a catch bond behavior for single αV-class integrins bound to vitronectin extends previous studies showing the same on fibronectin^37, 47^, and, thus, suggests catch bonds to be a more general feature amongst members of the integrin family. The higher rupture force in pKO-αV and pKO-αVβ5 fibroblasts and overall lower rupture force of pKO-αVβ3 fibroblasts recapitulated the overall behavior of cell adhesion force (Fig. 2b,d). However, the median rupture force of pKO-αV fibroblasts was intermediate to both pKO-αVβ3 and pKO-αVβ5 fibroblasts. This might be the case in pKO-αV fibroblasts due to the presence of both αVβ3 and αVβ5 integrins acting synergistically^51^. Thus, at the single integrin-vitronectin bond level, pKO-αV fibroblasts display tether forces and tether lifetimes similar to pKO-αVβ3 fibroblasts at lower retraction speeds (2 and 5 µm s^-1^). The non-linear rupture force in response to increasing retraction speed of pKO-αV fibroblasts is more similar to pKO-αVβ3 fibroblasts compared to pKO-αVβ5 fibroblasts, which indicates that the integrin β3-subunit dominates the β5-subunit if both are co-expressed. This domination of the integrin β3-subunit over the β5-subunit for the binding of vitronectin, is particularly interesting, since the β5-subunit complexed with the integrin αV-subunit establishes higher ligand-unbinding forces compared to αVβ3 integrin (Extended Data Fig. 5m,n). Thus, the integrin β5-subunit is the primary contributor toward establishing higher cell adhesion forces to vitronectin (Fig. 2h).

### Biphasic cell adhesion response requires actomyosin cortex regulation

To explore the roles of the intracellular components of the ECM-integrin-adaptor-cytoskeleton axis, we used small molecule inhibitors against components of the actomyosin cortex including F-actin (latrunculin A), myosin II (blebbistatin), Rho-associated coiled-coil containing protein kinase 1 (ROCK1, Y27632), and actin related protein 2/3 (Arp2/3) complex (CK666). First, we monitored for changes in cell surface expression levels of αVβ3 and αVβ5 integrins in response to compromised actomyosin and found them unaltered (Extended Data Fig. 6). Thereafter, we disrupted F-actin formation by latrunculin A that considerably reduced the cell adhesion force and led to a loss of the biphasic adhesion response in pKO-αV, pKO-αVβ3, and pKO-αVβ5 fibroblasts (Extended Data Fig. 7). Inhibition of myosin II and ROCK1 activity reduced the biphasic adhesion response at 5 µm s^-1^ whereas the adhesion forces for all other retraction speeds remained unaltered. Thus, ROCK1 inhibition prevented the biphasic adhesion response of pKO-αV and pKO-αVβ3 and pKO-αVβ5 fibroblasts. Furthermore, Arp2/3 inhibition disrupted the biphasic adhesion response for all three fibroblast lines (Fig. 3a). Although the adhesion force remained unchanged for pKO-αV fibroblasts, except at 5 µm s^-1^, the force increased for pKO-αVβ3 fibroblasts at higher retraction speeds while it reduced for pKO-αVβ5 fibroblasts. This opposite effect of Arp2/3 inhibition on pKO-αVβ3 and pKO-αVβ5 fibroblast adhesion indicates a differential regulation of Arp2/3 activity in an integrin β-subunit specific manner. In summary, an intact actin cortex is essential for adhesion initiation while responding to mechanical load, whereas the actomyosin contractility is required for the rapid increase of cell adhesion at low retraction speeds, which is the first phase of the biphasic cell adhesion response. Contrary to this finding, inhibition of the actin branches-inducing Arp2/3 complex triggered pKO-αVβ3 fibroblasts to establish higher adhesion force and pKO-αVβ5 fibroblasts to establish lower adhesion force at higher retraction speeds. This observation suggests that the Arp2/3 complex interacts differentially with integrin β3- and β5-subunits.

**Fig. 3:**
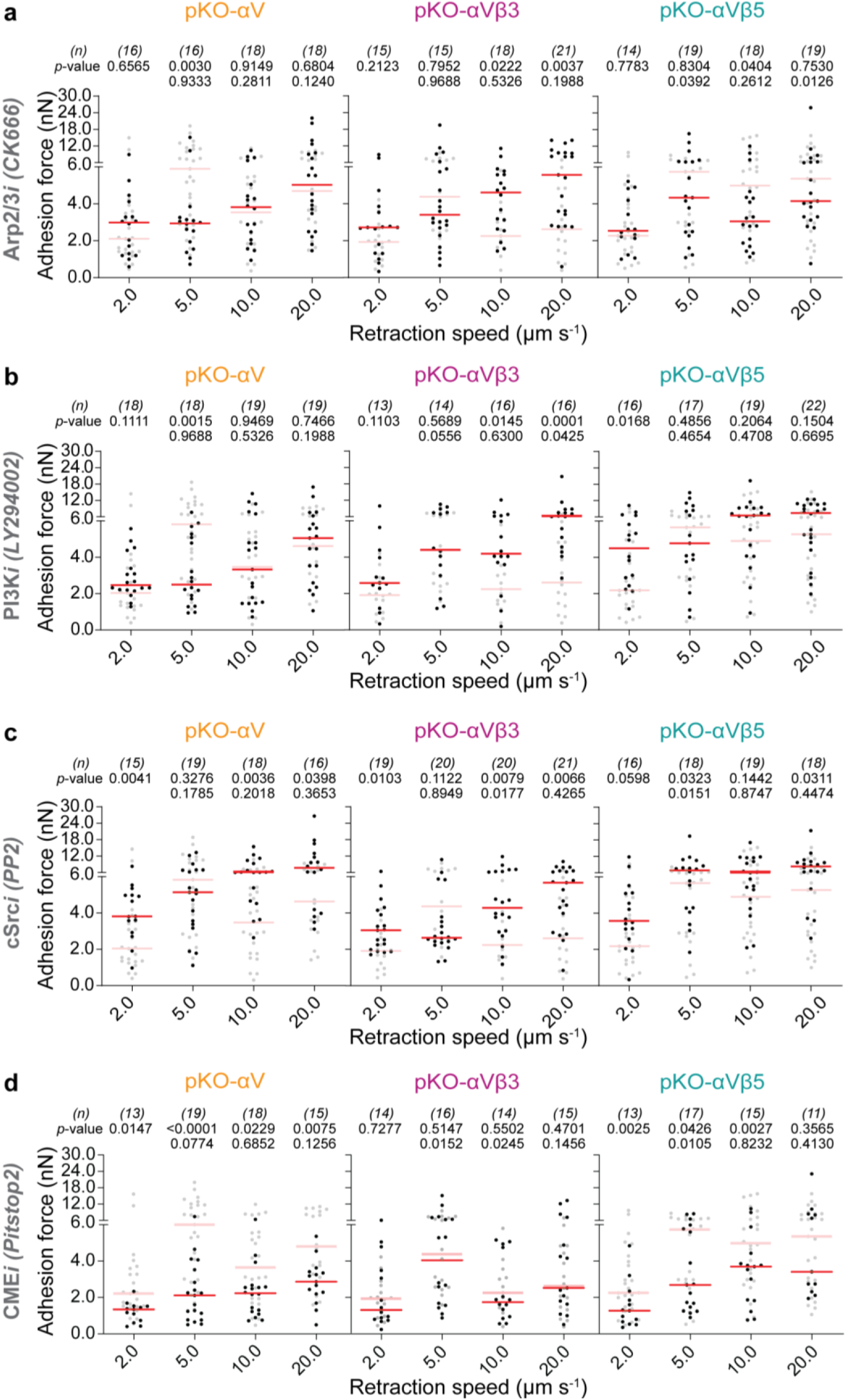
Arp2/3, PI3K, cSrc, and clathrin mediated endocytosis differentially mediate biphasic adhesion response to mechanical load in an integrin β-subunit specific manner. (**a**-**c**) Retraction speed-dependent adhesion force of chemically perturbed pKO-αV, pKO-αVβ3 and pKO-αVβ5 fibroblasts on vitronectin substrate after 120 s of contact. Fibroblasts were treated with (**a**) 20 μM Arp2/3 inhibitor CK666, (**b**) 10 μM phosphatidyl inositol 3-kinase (PI3K) inhibitor LY294002, (**c**) 20 μM cSrc kinase inhibitor PP2, and (**d**) 30 μM clathrin mediated endocytosis (CME) inhibitor Pitstop2. Each dot corresponds to one fibroblast, red bars to the median. *p*-values were obtained using two-tailed Mann-Whitney U-test. *n* denotes the number of cells measured. *p*-values in the top row compare the data to the unperturbed (lightened) data. *p*-values in the bottom row compare data for a given retraction speed to the preceding speed (e.g., 5 µm s^-1^ with respect to 2 µm s^-1^).

### β-subunit drives mechanotransduction of vitronectin-bound αV-class integrins

After characterizing the role of the actomyosin cortex in assisting the vitronectin-bound αV-class integrins in establishing the biphasic cell adhesion response, we explored the involvement of key signaling proteins including the focal adhesion kinase (FAK)^30, 52^, cellular Src kinase (cSrc)^24, 53^, and phosphatidyl inositol 3-kinase (PI3K). To perturb signaling proteins we used small molecules Y11 (inhibits FAK), PP2 (inhibits cSrc), or LY294002 (inhibits PI3K), which did not change the surface expression of αVβ3 or αVβ5 integrins (Extended Data Fig. 6). We observed that FAK inhibition reduced the adhesion force of pKO-αV and pKO-αVβ3 fibroblasts at 5 µm s^-1^, though their adhesion force at other retraction speeds remained unchanged similarly as of pKO-αVβ5 fibroblasts (Extended Data Fig. 7d). Thus, upon FAK inhibition, the integrin β3-subunit containing fibroblasts lost their ability to establish the biphasic adhesion response.

To test contributions of membrane phospholipids, we inhibited PI3K to prevent the formation of phosphatidyl inositol-3,4,5-trisposphate (PIP3) from phosphatidyl inositol-4,5-bisphosphate (PIP2) (Fig. 3b). In pKO-αV fibroblasts, PI3K-inhibition considerably reduced adhesion force at 5 µm s^-1^ retraction speed, but kept the adhesion force for 2, 10, and 20 µm s^-1^ unchanged. Thus, the biphasic adhesion response disappeared. pKO-αVβ3 fibroblasts responded to PI3K inhibition by reducing the biphasic adhesion response, however, the fibroblasts increased adhesion force considerably at 10 and 20 µm s^-1^. pKO-αVβ5 fibroblasts increased their adhesion force at 2 µm s^-1^, which disrupted the biphasic adhesion response, while the adhesion force at 5, 10, and 20 µm s^-1^ retraction speeds did not change significantly. Interestingly, cSrc inhibition disrupted the biphasic adhesion response of pKO-αV and pKO-αVβ3 fibroblasts, while pKO-αVβ5 fibroblasts maintained their biphasic adhesion response (Fig. 3c). These findings highlight that FAK is essential for the early cell adhesion response to mechanical load for both integrin β-subunits, while PI3K and cSrc signaling preferably influences αVβ3 integrins. Recent reports linked αVβ5 integrin to clathrin-mediated endocytosis (CME)^54^ and flat clathrin lattices^55^, we thus queried the involvement of this endocytosis in the early adhesion related biphasic response. Upon inhibiting CME by Pitstop2, the biphasic adhesion response was lost for αVβ5 integrin expressing pKO-αV and pKO-αVβ5 fibroblasts (Fig. 3d). On the contrary, pKO-αVβ3 fibroblasts maintained the biphasic response. Hence, the signaling pathways involved in the sensing of mechanical load through VN-bound αV-class integrins must differ for αVβ3 and αVβ5 integrins. To better understand the role of integrin β3/5-subunits in mechanotransduction, we transfected the pKO-αVβ3 fibroblasts with a chimeric β-subunit that contained the extracellular and transmembrane domain of the integrin β5-subunit fused to the cytoplasmic tail of the integrin β3-subunit. SCFS showed that pKO-αVβ3 fibroblasts transfected with this β-subunit chimera could not entirely recover the biphasic adhesion response such as observed for pKO-αV fibroblasts, unlike pKO-αVβ3 fibroblasts rescued with the full integrin β5-subunit (Fig. 1b and Extended Data Fig. 7). The biphasic adhesion response instead remained similar to pKO-αVβ3 fibroblasts (Fig. 2b). Thus, these data point to an involvement of the integrin β5-subunit tail, within seconds of binding to vitronectin, and hence implicate the co-existence of at least two different mechanosensing signal pathways depending on the β3- and β5-subunits.

### Vitronectin-bound αV-class integrins modulate cellular stiffness

The biphasic response for vitronectin-bound αV-class integrins emerged after 5 s of substrate contact in contrast to fibronectin-bound α5β1 integrins that trigger similar adhesion responses within 5 s^37, 47^. To further understand the role of αV-class integrins, we applied the Hertz-Sneddon model to extract the effective stiffness (Young’s modulus) of fibroblasts from SCFS data recorded at 120 s contact time (Fig. 4a). pKO-αV and pKO-αVβ5 fibroblasts increased stiffness contrary to pKO-αVβ3 fibroblasts with increasing retraction speed. These responses suggested the integrin β5-subunit may contribute to cellular stiffness regulation. To enquire whether expression of different integrin heterodimers and their respective mechanosensing lead to changes in overall cellular stiffness, we extracted the cellular stiffness from SCFS experiments for pKO-αV, pKO-αVβ3, and pKO-αVβ5 fibroblasts performed in the absence or presence of small molecule inhibitors against Arp2/3, myosin II, F-actin, PI3K, cSrc and CME (Fig. 4 and Extended Data Fig. 9). Inhibition of F-actin (Extended Data Fig. 9a), myosin II (Extended Data Fig. 9b), PI3K (Extended Data Fig. 9c), and cSrc (Extended Data Fig. 9d) reduced the stiffness of pKO-αV and pKO-αVβ5 fibroblasts for all retraction speeds, while pKO-αVβ3 fibroblasts increased stiffness mainly at higher retraction speeds for these inhibitions. Similarly, the inhibition of Arp2/3 or CME increased the stiffness of pKO-αVβ5 fibroblasts considerably (Fig. 4b,c). However, CME inhibition also increased the stiffness of pKO-αVβ3 fibroblasts considerably, whereas Arp2/3 inhibition showed no effect (Fig. 4c). Thus, independent of the integrin, CME might be involved in regulating the overall fibroblast stiffness. Moreover, the varied effect of Arp2/3 inhibition on the stiffness of pKO-αVβ3 and pKO-αVβ5 fibroblasts (Fig. 4b) suggests that Arp2/3, a known driver of CME, regulates CME only in the case of pKO-αVβ5 fibroblasts. We further characterized the αVβ5 selectivity of CME by quantifying clathrin light chain (CLC) assemblies in pKO-αV, pKO-αVβ3, and pKO-αVβ5 fibroblasts. Only the αVβ5 integrin expressing pKO-αV and pKO-αVβ5 fibroblasts formed clusters of CLC after 45 min of spreading on vitronectin (Fig. 4d). The area quantification of CLC assemblies showed that pKO-αV and pKO-αVβ5 fibroblasts form similar sized assemblies as opposed to pKO-αVβ3 fibroblasts (Fig. 4e). Thus, fibroblasts initiating adhesion to vitronectin regulate stiffness through integrin β-subunits and signaling inputs including those from CME cascade regulation.

**Fig. 4:**
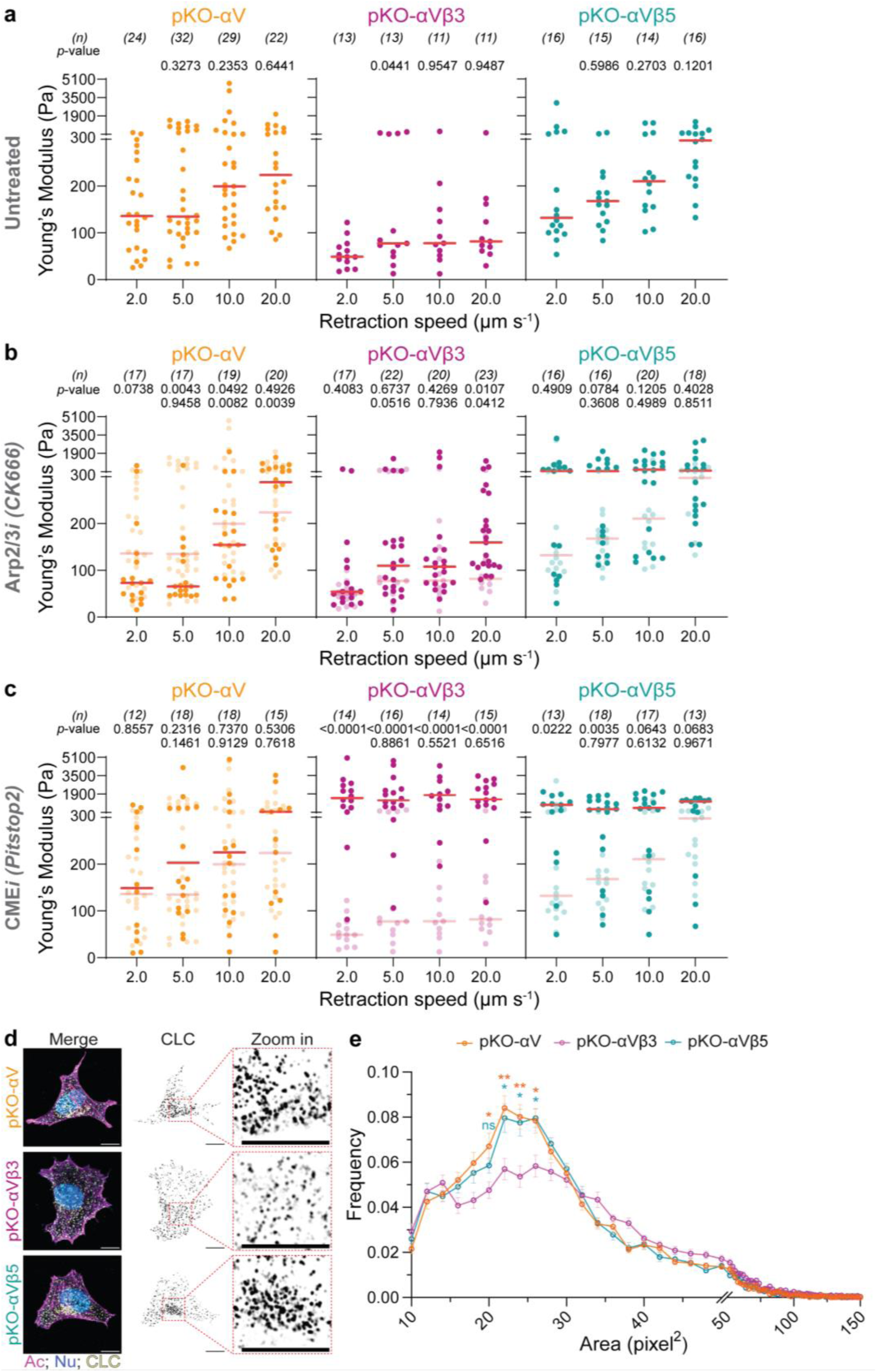
Integrin β-subunit identity, Arp2/3 and clathrin mediated endocytosis are key determinants of cytoskeletal stiffening under mechanical load. (**a-c**) Retraction speed-dependent apparent cell stiffness (Young’s modulus) determined for chemically (un-)/perturbed pKO-αV, pKO-αVβ3 and pKO-αVβ5 fibroblasts on vitronectin substrate after 120 s of contact. Fibroblasts were either (**a**) untreated or were treated with (**b**) 20 μM Arp2/3 inhibitor CK666, or (**c**) 30 μM clathrin mediated endocytosis (CME) inhibitor Pitstop2. Each dot represents one retraction curve corresponding to data reported in Fig. 2, B and C, and Fig. 3, A and D, red bars represent median values. *p*-values were obtained using two-tailed Mann-Whitney U-test. *n* denotes the number of cells measured. *p*-values in the top row compare the data to the unperturbed (lightened) data. *p*-values in the bottom row compare data for a given retraction speed to the preceding speed (e.g., 5 µm s^-1^ with respect to 2 µm s^-1^). **d**, Immunofluorescence images with labelled actin (Ac), nucleus (Nu) and clathrin light chain (CLC) in pKO-αV, pKO-αVβ3 and pKO-αVβ5 fibroblasts on vitronectin substrate after 45 min of adhesion. Zoom in highlighting the CLC assemblies in the respective fibroblasts. Scale bars, 10 μm. **e**, Frequency distribution of area quantified for CLC assemblies in pKO-αV, pKO-αVβ3 and pKO-αVβ5 fibroblasts. Each data point represents mean ± SE calculated for ≥15 fibroblasts. ‘ns’ non-significant; * *p*-values < 0.05; ** *p*-values < 0.01, obtained using Kruskal-Wallis/Dunn’s test. ‘Orange’ *p*-values represent comparison between pKO-αV and pKO-αVβ3 data. ‘Cyan’ *p*-values represent comparison between pKO-αVβ5 and pKO-αVβ3 data.

### αVβ3 integrin dominates long-term adhesome composition on vitronectin

To gain insight into the molecular underpinnings of the integrin β-subunit dependent biphasic response of fibroblast adhesion we quantitatively and qualitatively analyzed the adhesome composition. To this end, we cultured pKO-αV, pKO-αVβ3, and pKO-αVβ5 fibroblasts for 45 min on vitronectin- or poly-*L*-lysine-coated substrates, crosslinked and isolated the adhesome, released the crosslinks, trypsinized the proteins and analyzed them by mass spectrometry^27, 56^ (Extended Data Methods). Principal component analysis showed that the proteome of poly-*L*-lysine cultured fibroblasts clustered separately from vitronectin-cultured fibroblasts (Fig. 5a). Key components of the consensus adhesome like the adaptor proteins talin 1 (Tln1), kindlin 2 (Fermt2), vinculin (Vcl), zyxin (Zyx), integrin-linked pseudokinase (Ilk), parvin A (Parva) and paxillin (Pxn) as well as the signaling proteins C-terminal Src kinase (Csk), and focal adhesion kinase (Ptk2) exhibited no marked difference between pKO-αVand pKO-αVβ3 fibroblasts (Fig. 5a and Extended Data Fig. 10a,b). However, the consensus adhesome proteins associated preferentially to αVβ3 integrin (Fig. 5b,c), which indicates that αVβ3 integrin dominates the consensus adhesome assembly when both αVβ3 and αVβ5 integrins are present (Fig. 5c,d). Yet, we did not find adhesome components that have previously been associated with αVβ5 integrins in the context of long-term adhesion of human cancer cells^10^ or keratinocytes^57^, which is most likely due to the much longer adhesion time-frame and/or the different tissue origins. In summary, the data demonstrate a clear divergence between the adhesomes established through integrin β3- and β5-subunits of vitronectin-bound αV-class integrins and confirm that the β3-subunit takes a dominant role in assembling the long-term (45 min) adhesome in pKO-αV fibroblasts expressing both integrin β-subunits.

**Fig. 5:**
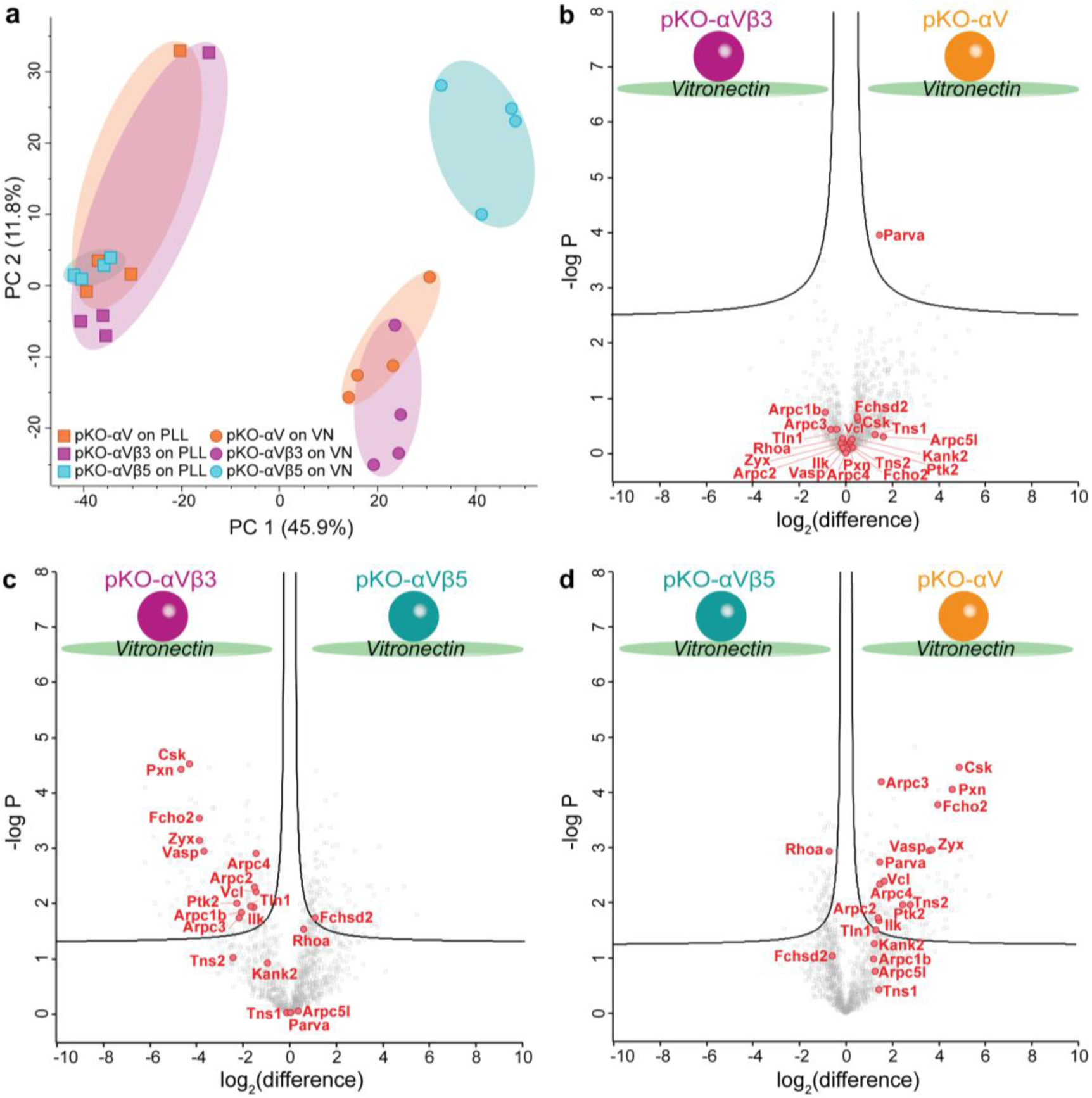
The consensus adhesome of αVβ3 integrin dominates the adhesome composition of pKO-αV fibroblasts on vitronectin. **a**, Principal component (PC) analysis for the mass spectrometry of fibroblast adhesome. Quality check for pKO-αV, pKO-αVβ3 and pKO-αVβ5 fibroblasts cultured on poly-*L*-lysine (squares) and vitronectin (circles) used for adhesome mass spectrometry analysis. PC 1 along the x-axis represents 45.9% variation based on the poly-*L*-lysine or vitronectin substrate the fibroblasts were cultured on. Four replicates for each condition are depicted. (**b-d**) Consensus adhesome components highlighted to display the comparison of relative protein abundance after 45 min of fibroblast adhesion to vitronectin for (**b**) pKO-αVβ3 and pKO-αV fibroblasts, (**c**) pKO-αVβ3 and pKO-αVβ5 fibroblasts, and (**d**) pKO-αVβ5 and pKO-αV fibroblasts, normalized to poly-L-lysine control. Black lines show the boundaries used to identify protein candidates corresponding to a false-discovery rate of 0.05 and a penalty factor for sample-wide standard deviation of 0.1. The data is representative of four independent replicates.

## Outlook

Integrin-mediated mechanotransduction has been studied substantially in recent years, especially the roles that individual β-subunits of the αV-class integrins play during substrate contextualization and establishment of adhesion complexes. For instance, αVβ3 integrins give rise to focal adhesions on fibronectin while αVβ5 integrins form focal and reticular adhesions, or flat clathrin lattices on vitronectin^55, 57-59^. Deciphering the molecular underpinnings of such integrin-mediated mechanotransduction is key to better understand cellular homeostasis and disease. We show that adhesomes of fibronectin or vitronectin adhered fibroblasts enrich for ECM proteins or complement system proteins, respectively (Fig. 1a). Thus, fibroblasts establish highly specialized and RGD-context dependent extracellular niches within tens of minutes. This finding aligns with previous reports indicating that fibronectin serves as a scaffold for collagen^60^, while vitronectin sequesters complement system proteins^61, 62^. αV-class integrins further enable rapid ECM sensing and faster spreading of fibroblasts on vitronectin than on fibronectin, thereby causing fibroblasts to occupy twice the area within 5 min (Fig. 1c,d). These findings highlight that αV-class integrins distinguish fibronectin from vitronectin and trigger substantially different cellular responses.

Based on previous studies exploring the role of varying mechanical load during early phases of cell adhesion^37, 41^, we hypothesize a similar occurrence within the αV-class integrins that might be crucial for mechanosensing fibronectin or vitronectin. Additionally, cells expressing αV-class integrins encounter mechanical cues ranging from intravascular pressure in the cardiovascular system to shear and strain in interstitial tissue^39, 63^. We show that within seconds of attaching to vitronectin, fibroblasts employ αV-class integrins to strengthen adhesion biphasically in response to increasing mechanical load (Fig. 1b). Fibroblasts perform this biphasic adhesion strengthening if adhering to vitronectin exclusively, but not to fibronectin (Fig. 1b and Extended Data Fig. 2). Both integrin β-subunits trigger the biphasic cell adhesion response differently at high (β5-subunit) or low (β3-subunit) adhesion force (Fig. 2b,d,g,i). We have reported a similar phenomenon for the α5β1 integrin-fibronectin bond supported by αVβ3 integrin^37^, since fibronectin is the cognate ligand for both integrins. Here, we add the vitronectin-binding of αVβ3 and αVβ5 integrin to the existing literature of the few integrins for which early adhesion and mechanosensation have been quantitatively described^37, 47^. Our findings further imply that mechanical forces, such as typically experienced during early phases of cell adhesion, enable integrin receptors to optimally bind and respond to their cognate ECM ligands. This ECM deciphering ability must further ensure physiological resilience for fibroblasts expressing αV-class integrins that encounter similar magnitudes of mechanical loads ranging from intravascular pressure in the cardiovascular system to shear and strain in interstitial tissue^39, 63^.

Next, we establish vitronectin as compliant extracellular ligand to which mechanically loaded fibroblasts respond biphasically upon early phases of engagement. Analysis of single integrin-vitronectin bonds shows that αVβ3 integrin dominates αVβ5 integrin in fibroblasts to establish adhesion to vitronectin. This dominance of αVβ3 integrin, which may originate from its higher intracellular engagement with talin and kindlin^51^ compared to αVβ5 integrin, leads to a higher likelihood of inside-out activation of αVβ3 integrin at the cell surface. Thus, despite having a higher affinity for binding to vitronectin, αVβ5 integrin is outcompeted by αVβ3 integrin, which likely has a higher avidity for binding to vitronectin, during the early phases of adhesion. Furthermore, the striking similarity between the αV-class integrin-vitronectin catch bond behavior and the biphasic cell adhesion response highlights that the catch bond phenomenon of single integrins is not merely mechanical, but also involves specific intracellular signaling pathways representing different phases of the catch bond. Thereby each compliant pathway is either triggered by αVβ5 or αVβ3 integrin.

The biphasic cell adhesion response facilitated by αV-class integrins requires force generation through actomyosin regulatory components like myosin II and ROCK1 (Extended Data Fig. 7a-c). Notably, Arp2/3 inhibition disrupts the β-subunit-dependent biphasic cell adhesion response to different degrees, suggesting that the Arp2/3 complex is differentially recruited to the integrin β3- and β5-subunit and regulates the adhesome within seconds (Fig. 3a). This additional role of Ap2/3 may arise from its involvement as a scaffold within the integrin adhesome, beyond its conventional function as an actin brancher^64^. We also show that FAK is crucial for the biphasic adhesion response triggered by both the integrin β3- and β5-subunits (Extended Data Fig. 7d), which extends previous studies in disease models^65, 66^ that showed the importance of active FAK for αVβ3- or αVβ5-integrin mediated angiogenesis and cancer progression, respectively. Although cSrc is reported to contribute to adhesion *via* β3- and β5-subunits in epithelial and cancer tissues^67-69^, we provide evidence that during early mechanosensing in fibroblasts, only the integrin β3-subunit requires active cSrc (Fig. 3c). In addition to the actomyosin regulation and adhesion related signaling, we discover that CME contributes to the αVβ5-integrin specific biphasic adhesion response of fibroblasts adhering to vitronectin within 120 s. We speculate that it is this early mechanosensory coupling and αVβ5-integrin specificity of clathrin that results in clathrin-linked adhesion complexes like clathrin-coated pits or flat clathrin lattices after minutes to hours of αVβ5-integrin mediated adhesion to vitronectin (Fig. 4d)^55, 70^. Thus, although the adhesome components, namely Arp2/3 (Fig. 3a), PI3K, cSrc and processes like CME contribute differentially to the integrin β-subunit-dependent biphasic cell adhesion response (Fig. 3b-d), we find components like F-actin, myosin II, ROCK1 (Extended Data Fig. 7a-c), and FAK (Extended Data Fig. 7d) taking a conserved roles for both β3- and β5-subunits.

Whether these proteins are also found in the integrin β-subunit associated adhesome at later adhesion times (>> 120 s) still remained unclear. Using mass spectrometry, we show that the integrin β3- and β5-subunits indeed assemble distinct adhesomes in fibroblast adhering for 45 min to vitronectin (Fig. 5). Additionally, pKO-αV fibroblasts containing both integrin β3-and β5-subunits show adhesome compositions similar to pKO-αVβ3 fibroblasts containing only the β3-subunit, which is unlike pKO-αVβ5 fibroblasts containing only the β5-subunit (Extended Data Fig. 10).

Moreover, our data suggests that the stiffness of fibroblast initiating adhesion to vitronectin is regulated through the integrin β-subunit (Fig. 4 and Extended Data Fig. 9). Thereby, αVβ5 integrin dominates over αVβ3 integrin in determining the cellular stiffness post adhesion within 120 s. This functional readout of rapid and early mechano-regulation of cell-scale biophysical properties by integrins also highlights the strength of our SCFS assay. Additionally, the integrin associated signaling pathways and actomyosin regulatory pathways also contribute to cellular stiffness within 120 s.

In summary, our study provides profound insights into how mammalian fibroblasts attaching to vitronectin establish and strengthen adhesion biphasically through αV-class integrins (Fig. 6a). We elucidate that these integrins individually mechanosense vitronectin and respond differentially to adhesion time and mechanical load. Within seconds of binding to vitronectin, the αV-class integrins trigger different intracellular signaling pathways (FAK, PI3K, cSrc, and CME) and regulatory components of the actomyosin cortex (myosin II, ROCK1, and Arp2/3) (Fig. 6b,c). These signaling pathways transduce a biphasic adhesion response that scales from the single integrin level to that of the entire fibroblast and in addition regulates cell mechanical properties (Fig. 4 and Extended Data Fig. 9). Of note is that αVβ3 integrin dominates the biphasic adhesion response, while αVβ5 integrin dominates in modulating the cellular biophysical properties like tether force and cell stiffness. At longer timescales of tens of minutes, the early (≤ 120 s) mechanosensitive assemblies mature into distinct integrin β-subunit-dependent adhesomes (Fig. 4d, Fig. 5, Fig. 6d and Extended Data Fig. 10), where molecules like Arp2/3, FAK, cSrc, Zyx, Vcl, Tln1, etc., enrich in an β3-subunit specific manner. Therefore, we comprehensively demonstrate that αVβ3 and αVβ5 integrin dominate distinct aspects of regulating the early (≤ 120 s) and late mechanotransduction of fibroblasts adhering to vitronectin, leading to highly customized cellular responses.

**Fig. 6:**
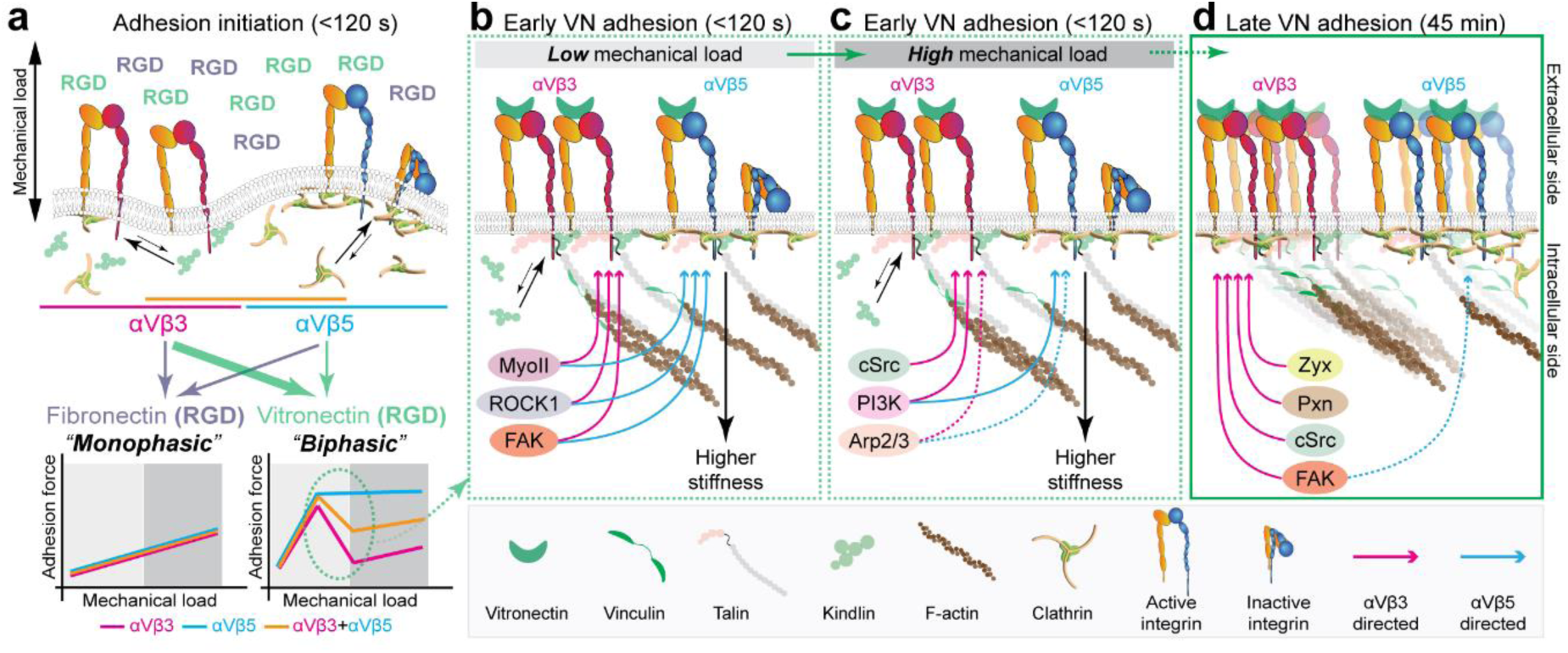
The β3- and β5-subunits of αV-class integrins contextualize the extracellular RGD-motif and take different roles in establishing cell adhesion and mechanics in response to mechanical load. **a**, Depending on which integrin β-subunit complexes with the αV integrin subunit, αVβ3 or αVβ5 integrins are formed, which both bind and contextualize the RGD-motif in fibronectin and vitronectin to initiate cell adhesion differentially. If the integrins bind the RGD-motif in fibronectin, “monophasic” adhesion strengthening occurs in response to mechanical load. However, if bound to the RGD-motif in vitronectin, fibroblasts strengthen adhesion in a “biphasic” manner in response to load. In this biphasic adhesion, αVβ3 integrin dominates over αVβ5 integrin, while αVβ5 integrin involves clathrin mediated endocytosis to contribute to cell adhesion strengthening and stiffening. **b** and **c**, Early mechanosensing within seconds, couples to mechanical load. **b,** At low mechanical load, αVβ3 and αVβ5 integrins strengthen cell adhesion to vitronectin (VN) by involving intracellular signaling molecules and adhesome proteins like MyoII, ROCK1 and FAK. **c**, With increasing mechanical load, αV-class integrins recruit different molecules like cSrc, PI3K, and Arp2/3 to strengthen adhesion (dotted lines). Here, αVβ3 integrin is the main regulator of signaling pathways, while αVβ5 integrin is the main regulator of higher cell stiffness. **d**, Late adhesome composition for cell adhesion to vitronectin is distinct for αVβ3 and αVβ5 integrins, where αVβ3 integrin dominates over αVβ5 integrin by recruiting most consensus adhesome components, including Zyx, Pxn, cSrc, and FAK, in addition to the proteins described in early adhesion formation (**b**, **c**).

## References

1. Swaminathan, V. et al. Actin retrograde flow actively aligns and orients ligand-engaged integrins in focal adhesions. Proceedings of the National Academy of Sciences 114, 10648–10653 (2017).

2. Plotnikov, Sergey V., Pasapera, Ana M., Sabass, B. & Waterman, Clare M. Force Fluctuations within Focal Adhesions Mediate ECM-Rigidity Sensing to Guide Directed Cell Migration. Cell 151, 1513–1527 (2012).

3. Schwarz, U.S. & Gardel, M.L. United we stand – integrating the actin cytoskeleton and cell–matrix adhesions in cellular mechanotransduction. Journal of Cell Science 125, 3051–3060 (2012).

4. Riveline, D. et al. Focal Contacts as Mechanosensors: Externally Applied Local Mechanical Force Induces Growth of Focal Contacts by an Mdia1-Dependent and Rock-Independent Mechanism. Journal of Cell Biology 153, 1175–1186 (2001).

5. Harburger, D.S. & Calderwood, D.A. Integrin signalling at a glance. Journal of Cell Science 122, 1472–1472 (2009).

6. Moser, M., Legate, K.R., Zent, R. & Fässler, R. The Tail of Integrins, Talin, and Kindlins. Science 324, 895–899 (2009).

7. Chastney, M.R. et al. Topological features of integrin adhesion complexes revealed by multiplexed proximity biotinylation. Journal of Cell Biology 219 (2020).

8. Horton, E.R. et al. Definition of a consensus integrin adhesome and its dynamics during adhesion complex assembly and disassembly. Nature Cell Biology 17, 1577–1587 (2015).

9. Byron, A. et al. A proteomic approach reveals integrin activation state-dependent control of microtubule cortical targeting. Nature Communications 6, 6135 (2015).

10. Paradžik, M. et al. KANK2 Links αVβ5 Focal Adhesions to Microtubules and Regulates Sensitivity to Microtubule Poisons and Cell Migration. Frontiers in Cell and Developmental Biology 8, 1–17 (2020).

11. Kanchanawong, P. & Calderwood, D.A. Organization, dynamics and mechanoregulation of integrin-mediated cell–ECM adhesions. Nature Reviews Molecular Cell Biology 24, 142–161 (2023).

12. Piccolo, S., Sladitschek-Martens, H.L. & Cordenonsi, M. Mechanosignaling in vertebrate development. Developmental Biology 488, 54–67 (2022).

13. Sun, Z., Costell, M. & Fässler, R. Integrin activation by talin, kindlin and mechanical forces. Nature Cell Biology 21, 25–31 (2019).

14. Sun, Z., Guo, S.S. & Fässler, R. Integrin-mediated mechanotransduction. Journal of Cell Biology 215, 445–456 (2016).

15. Hashiguchi, S., Tanaka, T., Mano, R., Kondo, S. & Kodama, S. CCN2-induced lymphangiogenesis is mediated by the integrin αvβ5–ERK pathway and regulated by DUSP6. Scientific Reports 12, 926 (2022).

16. Oguri, Y. et al. CD81 Controls Beige Fat Progenitor Cell Growth and Energy Balance via FAK Signaling. Cell 182, 563–577 e520 (2020).

17. Henderson, N.C. et al. Targeting of αv integrin identifies a core molecular pathway that regulates fibrosis in several organs. Nature Medicine 19, 1617–1624 (2013).

18. Desgrosellier, Jay S. et al. Integrin αvβ3 Drives Slug Activation and Stemness in the Pregnant and Neoplastic Mammary Gland. Developmental Cell 30, 295–308 (2014).

19. Hatley, R.J.D. et al. An αv-RGD Integrin Inhibitor Toolbox: Drug Discovery Insight, Challenges and Opportunities. Angewandte Chemie International Edition 57, 3298–3321 (2018).

20. Hurtado de Mendoza, T., et al. Tumor-penetrating therapy for β5 integrin-rich pancreas cancer. Nature Communications 12, 1541 (2021).

21. Pearson, J.D. et al. Binary pan-cancer classes with distinct vulnerabilities defined by pro- or anti-cancer YAP/TEAD activity. Cancer Cell 39, 1115–1134.e1112 (2021).

22. Rahman, S.R. et al. Integrins as a drug target in liver fibrosis. Liver International 42, 507–521 (2022).

23. Slack, R.J., Macdonald, S.J.F., Roper, J.A., Jenkins, R.G. & Hatley, R.J.D. Emerging therapeutic opportunities for integrin inhibitors. Nature Reviews Drug Discovery 21, 60–78 (2022).

24. Desgrosellier, J.S. et al. An integrin αvβ3–c-Src oncogenic unit promotes anchorage-independence and tumor progression. Nature Medicine 15, 1163–1169 (2009).

25. Balaban, N.Q. et al. Force and focal adhesion assembly: a close relationship studied using elastic micropatterned substrates. Nature Cell Biology 3, 466–472 (2001).

26. Sun, Z. Nascent Adhesions: From Fluctuations to a Hierarchical Organization. Current Biology 24, R801–R803 (2014).

27. Schiller, H.B. et al. β1- and αv-class integrins cooperate to regulate myosin II during rigidity sensing of fibronectin-based microenvironments. Nature Cell Biology 15, 625–636 (2013).

28. Gardel, M.L. et al. Traction stress in focal adhesions correlates biphasically with actin retrograde flow speed. Journal of Cell Biology 183, 999–1005 (2008).

29. Saraswathibhatla, A., Indana, D. & Chaudhuri, O. Cell–extracellular matrix mechanotransduction in 3D. Nature Reviews Molecular Cell Biology 24, 495–516 (2023).

30. Zhou, D.W. et al. Force-FAK signaling coupling at individual focal adhesions coordinates mechanosensing and microtissue repair. Nature Communications 12, 2359 (2021).

31. Brakebusch, C. & Fässler, R. The integrin–actin connection, an eternal love affair. The EMBO Journal 22, 2324–2333 (2013).

32. Chen, N.-P., Sun, Z. & Fässler, R. The Kank family proteins in adhesion dynamics. Current Opinion in Cell Biology 54, 130–136 (2018).

33. Schiller, H.B. & Fässler, R. Mechanosensitivity and compositional dynamics of cell–matrix adhesions. EMBO reports 14, 509–519 (2013).

34. Bachmann, M. et al. Induction of ligand promiscuity of αVβ3 integrin by mechanical force. Journal of Cell Science 133, jcs242404 (2020).

35. Grudtsyna, V., Packirisamy, S., Bidone, T.C. & Swaminathan, V. Extracellular matrix sensing via modulation of orientational order of integrins and F-actin in focal adhesions. Life Science Alliance 6, e202301898 (2023).

36. Polacheck, W.J. & Chen, C.S. Measuring cell-generated forces: a guide to the available tools. Nature Methods 13, 415–423 (2016).

37. Strohmeyer, N., Bharadwaj, M., Costell, M., Fässler, R. & Müller, D.J. Fibronectin-bound α5β1 integrins sense load and signal to reinforce adhesion in less than a second. Nature Materials 16, 1262–1270 (2017).

38. Lin, G.L. et al. Activation of beta 1 but not beta 3 integrin increases cell traction forces. FEBS Letters 587, 763–769 (2013).

39. Moore, S.W., Roca-Cusachs, P. & Sheetz, M.P. Stretchy Proteins on Stretchy Substrates: The Important Elements of Integrin-Mediated Rigidity Sensing. Developmental Cell 19, 194–206 (2010).

40. Sheetz, M. A Tale of Two States: Normal and Transformed, With and Without Rigidity Sensing. Annual Review of Cell and Developmental Biology 35, 169–190 (2019).

41. Elosegui-Artola, A. et al. Mechanical regulation of a molecular clutch defines force transmission and transduction in response to matrix rigidity. Nature Cell Biology 18, 540–548 (2016).

42. Jo, M.H. et al. Single-molecule characterization of subtype-specific β1 integrin mechanics. Nature Communications 13, 7471 (2022).

43. Benoit, M., Gabriel, D., Gerisch, G. & Gaub, H.E. Discrete interactions in cell adhesion measured by single-molecule force spectroscopy. Nature Cell Biology 2, 313–317 (2000).

44. Friedrichs, J., Helenius, J. & Muller, D.J. Quantifying cellular adhesion to extracellular matrix components by single-cell force spectroscopy. Nature Protocols 5, 1353–1361 (2010).

45. Krieg, M. et al. Atomic force microscopy-based mechanobiology. Nature Reviews Physics 1, 41–57 (2019).

46. Viljoen, A. et al. Force spectroscopy of single cells using atomic force microscopy. Nature Reviews Methods Primers 1, 63 (2021).

47. Andreu, I. et al. The force loading rate drives cell mechanosensing through both reinforcement and cytoskeletal softening. Nature Communications 12, 4229 (2021).

48. Polacheck, W.J., Charest, J.L. & Kamm, R.D. Interstitial flow influences direction of tumor cell migration through competing mechanisms. Proceedings of the National Academy of Sciences 108, 11115–11120 (2011).

49. te Riet, J., et al. Dynamic coupling of ALCAM to the actin cortex strengthens cell adhesion to CD6. Journal of Cell Science 127, 1595–1606 (2014).

50. Krieg, M., Helenius, J., Heisenberg, C.-P. & Muller, D.J. A Bond for a Lifetime: Employing Membrane Nanotubes from Living Cells to Determine Receptor–Ligand Kinetics. Angewandte Chemie International Edition 47, 9775–9777 (2008).

51. Sun, Z. et al. Kank2 activates talin, reduces force transduction across integrins and induces central adhesion formation. Nature Cell Biology 18, 941–953 (2016).

52. Cheng, B. et al. Nanoscale integrin cluster dynamics controls cellular mechanosensing via FAKY397 phosphorylation. Science Advances 6, eaax1909 (2020).

53. De Mets, R. et al. Cellular tension encodes local Src-dependent differential β1 and β3 integrin mobility. Molecular Biology of the Cell 30, 181–190 (2018).

54. Alfonzo-Méndez, M.A., Sochacki, K.A., Strub, M.-P. & Taraska, J.W. Dual clathrin and integrin signaling systems regulate growth factor receptor activation. Nature Communications 13, 905 (2022).

55. Zuidema, A. et al. Mechanisms of integrin αVβ5 clustering in flat clathrin lattices. Journal of Cell Science 131, jcs221317 (2018).

56. Schiller, H.B., Friedel, C.C., Boulegue, C. & Fässler, R. Quantitative proteomics of the integrin adhesome show a myosin II-dependent recruitment of LIM domain proteins. EMBO reports 12, 259–266 (2011).

57. Zuidema, A. et al. Molecular determinants of αVβ5 localization in flat clathrin lattices: Role of αVβ5 in cell adhesion and proliferation. Journal of Cell Science 135, jcs.259465 (2022).

58. Baschieri, F. et al. Frustrated endocytosis controls contractility-independent mechanotransduction at clathrin-coated structures. Nature Communications 9, 3825 (2018).

59. Elkhatib, N. et al. Tubular clathrin/AP-2 lattices pinch collagen fibers to support 3D cell migration. Science 356, eaal4713 (2017).

60. Saunders, J.T. & Schwarzbauer, J.E. Fibronectin matrix as a scaffold for procollagen proteinase binding and collagen processing. Molecular Biology of the Cell 30, 2218–2226 (2019).

61. Milis, L., Morris, C.A., Sheehan, M.C., Charlesworth, J.A. & Pussell, B.A. Vitronectin-mediated inhibition of complement: Evidence for different binding sites for C5b-7 and C9. Clinical and Experimental Immunology 92, 114–119 (1993).

62. Preissner, K.T. The role of vitronectin as multifunctional regulator in the hemostatic and immune systems. Blut 59, 419–431 (1989).

63. Chaudhuri, O., Cooper-White, J., Janmey, P.A., Mooney, D.J. & Shenoy, V.B. Effects of extracellular matrix viscoelasticity on cellular behaviour. Nature 584, 535–546 (2020).

64. Chorev, D.S., Moscovitz, O., Geiger, B. & Sharon, M. Regulation of focal adhesion formation by a vinculin-Arp2/3 hybrid complex. Nature Communications 5, 3758 (2014).

65. Cai, W.-J. et al. Activation of the integrins α5β1 and αvβ3 and focal adhesion kinase (FAK) during arteriogenesis. Molecular and Cellular Biochemistry 322, 161–169 (2009).

66. Tancioni, I. et al. FAK Inhibition Disrupts a β5 Integrin Signaling Axis Controlling Anchorage-Independent Ovarian Carcinoma Growth. Molecular Cancer Therapeutics 13, 2050–2061 (2014).

67. Eliceiri, B.P. et al. Src-mediated coupling of focal adhesion kinase to integrin αvβ5 in vascular endothelial growth factor signaling. Journal of Cell Biology 157, 149–160 (2002).

68. Arias-Salgado, E.G. et al. Src kinase activation by direct interaction with the integrin β cytoplasmic domain. Proceedings of the National Academy of Sciences 100, 13298–13302 (2003).

69. Bianchi-Smiraglia, A. et al. Integrin-β5 and zyxin mediate formation of ventral stress fibers in response to transforming growth factor β. Cell Cycle 12, 3377–3389 (2013).

70. Lock, J.G. et al. Clathrin-containing adhesion complexes. Journal of Cell Biology 218, 2086–2095 (2019).

