## Supplementary material for "Load-dependent RGD-context sensing *via* αV-class integrin reprograms cellular adhesion and mechanics within seconds": Supplemetary Material

**Methods**

Cell culture

All mouse embryonic fibroblast cell lines were cultured in Dulbecco modified eagle medium (DMEM, 31966047, Thermo Fisher Scientific), supplemented with 10% (v/v) fetal bovine serum (FBS, F9665, Sigma), 100 U $ml^{-1}$ penicillin and 100 µg $ml^{-1}$ streptomycin (15140122, Thermo Fisher Scientific). Fibroblasts were grown in tissue culture flasks (TCF012050, Jet BioFil) that were coated with bovine fibronectin (341631, Calbiochem, Merck Millipore). All fibroblast lines were routinely tested for mycoplasma contamination (MycoTest Kit, Merck) and cleaned, if required (MycoClear Kit, Merck).

Fibroblast engineering

To deplete β3/β5 integrins in pKO-αV fibroblasts, CRISPR/Cas9 was employed as described[1]. Briefly, single-guide RNAs (sgRNAs) for two different exons were designed using CHOPCHOP (https://chopchop.cbu.uib.no)[2] for *Itgb3* and *Itgb5*. genes in *Mus musculus* genome (mm10/GRCm38). sgRNAs were selected based on high rank, low self-complementarity, low off-target score, and efficiency score. The sgRNAs (with underlined PAM sequences; 5'-NGG-3') targeted two separate exons [*Itgb3* exon 4 (chr11:104,633,595-104,633,614) 5'-ATGTACATGTACGGCGATACAGG-3'; *Itgb3* exon 10 (chr11:104,643,724-104,643,743) 5'-CCAGCCCACGCTGCAACAATGGG-3'; Itgb5 exon 3 (chr16:33,865,447-33,865,466) 5'-ACCGTGGATTGCCAAAGTACTGG-3'; Itgb5 exon 4 (chr16:33,876,034-33,876,053) 5'-ACCCAATACACGGATTGGTCTGG -3'] on these genes. The forward and reverse DNA oligomers encoding sgRNAs were phosphorylated using T4 polynucleotide kinase enzyme (New England BioLabs, M0201S) and then hybridized. Following a BpiI (Thermo Fisher Scientific, ER1011) mediated digestion of backbones containing Cas9 and fluorescent reporter protein expression cassettes (pSpCas9-2A-BFP and pSpCas9-2A-GFP), sgRNA hybrids were ligated to the linearized backbones using T4 DNA ligase (New England BioLabs, M0202S). pKO-αV fibroblasts were then transfected using lipofectamine 3000 (Invitrogen) following the product manual. After 48 h of transfection, green fluorescence protein (GFP) and blue fluorescence protein (BFP) double positive fibroblasts were sorted (SONY MA900) and cultured on fibronectin-coated multi-well culture dishes. Loss of respective integrin expression was verified using immunolabelling and flowcytometric analysis (see Methods "*Flow cytometry*" section). To reintroduce integrins in KO fibroblasts, we cultured fibroblasts on fibronectin-coated 6 well plate. After attaining a confluency of ~70 %, we transfected the fibroblasts with a pCMV3-mItgb3 plasmid (Sino Biological Inc., MG50080-UT) and pCMV-mItgb5 plasmid (Sino Biological Inc., MG58180-CM) using Lipofectamine 3000. The transfection medium was washed off after ~ 20 h and replaced with culture medium containing 50 μg $ml^{-1}$ hygromycin B (Millipore, 400053). The hygromycin selection was continued for 3 days with daily replenishment of new antibiotic containing culture medium. After 3 days of selection, the surviving fibroblasts were stained with anti-mouse CD61 (integrin β3) (1:200, Invitrogen, #12-0611-82) with Armenian hamster IgG control (1:200, Invitrogen, #12-4888-83), anti-αVβ5 (1:200, BD Pharmigen, #565836) with mouse IgGb2(κ) control (1:200, BD Pharmigen, #565378). for flow cytometric analysis (see Materials and Methods "Flow cytometry experiments" section). Only cells with positive signal comparable to the parental fibroblasts were sorted using Sony MA900, expanded, and further used for SCFS, cell size, and integrin expression measurements.

Coating of AFM cantilever and Petri dishes

200 μm long tipless V-shaped silicon nitride cantilevers (NP-O: D, Bruker Inc.) were functionalized as described [3]. Briefly, cantilevers were plasma cleaned (PDC-32G, Harrick Plasma) for 2-3 minutes and incubated overnight with 2 mg $ml^{-1}$ concanavalin A (ConA, C2010, Sigma-Aldrich) at 4 °C. The fibronectin fragment (FNIII7-10) and RGD-deleted fibronectin fragment (FNIII7-10ΔRGD) used as substrates were expressed and purified as described before [4, 5]. For substrate coating, four-segmented polydimethylsilane (PDMS, 101697, Dowsil) masks were attached to the glass surface of a Petri dish (FD35-100, WPI). The PDMS-free windows of the masks were incubated with 20 μl of vitronectin, fibronectin, or FNIII7-10ΔRGD (all in 50 μg $ml^{-1}$ in PBS) overnight at 4 °C.

Single-cell force spectroscopy (SCFS)

All fibroblasts cell lines were grown on fibronectin-coated 24-well plates to a maximal confluency of ~80% (142475, Thermo Fisher Scientific) and serum-deprived in FBS-free DMEM with 100 U $ml^{-1}$ penicillin and 100 μg $ml^{-1}$ streptomycin (15140122, Thermo Fisher Scientific) overnight before the experiment. SCFS was performed using an AFM (CellHesion 200, JPK Instruments) and a motorized stage (JPK Instruments) mounted on an inverted optical microscope (AxioObserver Z1, Zeiss). The temperature during the measurements was maintained at 37°C by a temperature controlling system (The Cube, Life Imaging Services). Functionalized 200 μm long tipless V shaped silicon nitride cantilevers (NP-O: D, Bruker) with a nominal spring constant of 0.06 N $m^{-1}$ were used.

Fibroblasts were detached from the fibronectin-coated culture using 0.25% trypsin (Thermo) for 2 min. Trypsin was inactivated by resuspending the cells in SCFS medium [(DMEM, 12800017, Thermo Fisher) with 20 mM HEPES (A3724, Applichem) and 4 mM sodium bicarbonate, pH 7.2)] with added 1% (v/v) FBS, then the cells were pelleted and resuspended in 100 μl SCFS medium. Fibroblasts were incubated at 37°C to recover from 0.25% trypsin treatment for 30 min in SCFS medium. In the meantime, ECM protein functionalized glass-bottom Petri dishes were washed twice with 1X PBS to remove unbound ECM proteins. Next, suspended fibroblasts were added onto FNIII7-10ΔRGD-coated window in the glass-bottom dish. The ConA-coated cantilever was lowered onto a single fibroblast with an approach speed of 10 $\mu m\ s^{-1}$ until a contact force of 5 nN was attained. After a contact time of 5 s, the cantilever was retracted with 10 $\mu m\ s^{-1}$ until the fibroblast entirely detached from the substrate. The morphological state of the fibroblasts was continuously monitored throughout the experiment using an optical microscope. Cell adhesion measurements were conducted exclusively on non-blebbing, round, and non-spread fibroblasts of similar diameter. The adhesion force of each fibroblast was quantified by lowering the cantilever onto the substrate-coated glass dish with a speed of 5 $\mu m\ s^{-1}$. After a contact force of 1 nN was reached, the cantilever height was maintained for contact times of 5, 20, 50 or 120 s. Afterwards, the cantilever was retracted with 2, 3, 5, 6, 8, 10, 20 or 40 $\mu m\ s^{-1}$ for 100 μm until fibroblast was entirely separated from the substrate (Extended Data Fig. 1a). Adhesion forces were determined after the retraction force-distance curves were drift- and baseline-corrected using the JPK data processing software (version 8.0.92, Bruker Nano GmbH).

#### Adhesion probability assay

To perform the adhesion probability assays with single molecule sensitivity, the AFM (NanoWizard II, JPK Instruments) was mounted on an inverted optical microscope (AxioObserver Z1, Zeiss). The temperature of the experimental setup was maintained at 37 °C by a PetriDishHeater (JPK Instruments). Before the measurement, the glass-bottom Petri dish was coated with 50 $\mu g\ ml^{-1}$ of vitronectin or FNIII7-10ΔRGD substrate for 2 h at room temperature and was washed thrice with 1X PBS. Fibroblasts were attached to the ConA-coated cantilever as described, before being approached and retracted repeatedly from the substrate coated Petri dish. The cantilever-attached fibroblast was approached with 1 $\mu m\ s^{-1}$ speed to the substrate-coated support until reaching a contact force of 500 pN. The cantilever was retracted immediately afterwards, resulting in a contact time of ~50 ms between fibroblast and substrate. To estimate the binding probability of ligand-receptor bonds, every force-distance curve was analyzed to count the number of binding events using the data processing software of the AFM ((version 8.0.92, Bruker Nano GmbH); Extended Data Fig. 1c). The adhesion probability was calculated for each fibroblast as the number of force-distance curves showing single adhesion events over the total number of curves recorded.

#### Analysis of single rupture and tether events

Tether and rupture forces were extracted from force-distance curves using in-house code (Igor 8, Wavemetrics). To analyse single integrin bonds, force-distance curves, acquired as described in the 'SCFS' section, for pKO-αV, pKO-αVβ3, and pKO-αVβ5 fibroblasts. Rupture events were analysed to measure the forces rupturing single integrin-vitronectin bonds. Tether events represent the force required to mechanically extract a membrane tether from the cell [6]. Tether lifetimes were calculated by dividing tether length with the retraction speed during detachment of fibroblasts from vitronectin. The selection of tether and rupture events was done based on previous reports.

#### Analysis of apparent cell stiffness

We used JPK data processing software (version 8.0.92, Bruker Nano GmbH) to drift- and baseline-correct the force-distance curves. The curves were smoothened using Boxcar method with a width of 3.0. The retraction curves were then fitted between the values of zero force and the maximum force before cell detachment to obtain the apparent cell stiffness (Young's modulus). The fit was performed using the Hertz-Sneddon model for a spherical tip shape of tip radius 10 μm and Poisson ratio of 0.5.

Perturbation

Cytoplasmic proteins were perturbed by incubating fibroblasts at 37°C for 30 min in SCFS media with inhibitors as follows; 1 μM latrunculin A (LatA, Sigma), 200 μM CK666 (Tocris Bioscience), 20 μM blebbistatin (Sigma), 10 μM Y27632 (Sigma), 10 μM Y11 (Tocris Bioscience), 10 μM LY249002 (Cell Signaling Technology), or 20 μM PP2 (Tocris Bioscience) in DMSO (to a final composition of 0.1% v/v DMSO in SCFS medium). All inhibitors were present in the given concentrations during SCFS. 0.1% v/v DMSO was tested as solvent control to confirm no effect on adhesion forces from carrier solvents.

Flow cytometry experiments

For flow cytometry analysis, fibroblasts were cultured on fibronectin-coated 24 well plates (Greiner) up to ≈ 80% confluency. They were serum deprived for > 4 h in DMEM (31966047, Thermo Fisher Scientific), supplemented with 100 U $ml^{-1}$ penicillin and 100 μg $ml^{-1}$ streptomycin (15140122, Thermo Fisher Scientific). Serum-deprived fibroblasts were detached from the plates using 0.25% (w/v) trypsin-EDTA (Gibco #25200056) at 37°C for 2 min. Detached fibroblasts were resuspended in SCFS media with 1% (v/v) FBS and recovered from detachment process at 37°C for at least 30 min. After recovery, cells were pelleted and $10^5$ fibroblasts were resuspended in 100 µl of flow cytometry buffer (2 mM EDTA and 3% (w/v) BSA in PBS) containing fluorophore conjugated primary antibodies as follows; anti-mouse CD61 (integrin β3) (1:200, Invitrogen, #12-0611-82) with Armenian hamster IgG control (1:200, Invitrogen, #12-4888-83), anti-αVβ5 (1:200, BD Pharmigen, #565836) with mouse IgGb2(κ) control (1:200, BD Pharmigen, #565378). The cells were stained with the said antibodies for 30 mins at 4°C in dark conditions. Afterwards, fibroblasts were washed using flow cytometry buffer twice, followed by measuring their fluorescence intensity employing LSRFortessa (BD AG). The flow cytometry data was analyzed by using FlowJo (v10, BD AG). Fluorescence activated cell sorting (FACS) was performed using the culture and staining protocols identical to the flow cytometry analysis experiments. Integrin-negative cells were sorted using a cell sorter (MA900 Sony) at 4°C in fibronectin-coated 6-well plates.

Cell spreading kinetics

To monitor fibroblasts spreading on vitronectin and fibronectin, we used an airyscan confocal microscope (LSM980, Zeiss), a 20x air objective (Zeiss), and data acquisition software (ZEN Blue, Zeiss). pKO-αV fibroblasts were stained with CellTracker™ Green CMFDA (5-chloromethylfluorescein diacetate) dye (excitation/emission spectra of 492/517 nm maxima) (Thermo Fisher Scientific #C7025) to stain the cytoplasm. The fibroblasts were added to fibronectin- and vitronectin-coated Ibidi well plates with inserts and monitored for their spreading every 30 s using confocal microscopy for > 25 min. Laser intensities and gains were kept constant across the experiments. Fluorescence signal was analyzed using Fiji (Version 2.1.0/1.52c). White outlines masks used to estimate the projected cell areas were obtained using suitable threshold settings and automated detection of the region of interest around the CellTracker™ signal.

Adhesome proteomics

Quantification of adhesome composition was performed by enrichment and isolation of the adhesion associated proteome, as described (*29, 59*). Briefly, for each fibroblast line four 10 cm diameter Petri dishes per experimental condition were coated with 3 ml of 50 μg $ml^{-1}$ fibronectin, 50 μg $ml^{-1}$ vitronectin, or 0.1% v/v poly-*L*-lysine overnight at 4 °C. 70-90% confluent pKO-αV, pKO-αVβ3, and pKO-αVβ5 fibroblasts were serum-starved for 4 h before being detached with 0.008% (v/v) trypsin-EDTA , obtained by diluting 0.25% (w/v) trypsin-EDTA (Gibco #25200056) in 1X PBS, for 3-5 min, neutralized using 10% (v/v) trypsin inhibitor (in FBS-free DMEM), and allowed to recover from detachment in FBS-free DMEM for 30 min at 37 °C. The fibroblasts were plated on the substrate-coated dishes for 45 min. Afterwards, following a wash with 1X PBS, each dish was incubated on ice with 5 ml PBS (supplemented with 1 mM $CaCl_2$ and 1 mM $MgCl_2$). Next, samples were treated with 5 ml crosslinker solution (25 mM dithiobis succinimidyl propionate (DSP, 22585, Sigma) and 25 mM 1,4-di-[3′-(2′-pyridyldithio)-propionamido] butane (DPDPB, CL112, Soltec Ventures) in PBS supplemented with 1 mM $CaCl_2$ and 1 mM $MgCl_2$) for 5 min at room temperature on a rocker. The crosslinking reaction was quenched for 15 min at room temperature by adding 100 mM Tris (pH 7.5) to each dish. Fibroblasts were lysed for 30 min at 4 °C in 3 ml radioimmunoprecipitation assay (RIPA) lysis buffer (25 mM Tris, 150 mM NaCl, 0.1% (w/v) SDS, 0.5% (w/v) sodium deoxycholic acid, one protease inhibitor tablet per 50 ml of RIPA lysis buffer (04693159001 Roche, Sigma)). Non-covalently bound cellular material was removed by applying high-pressure shear-flow jetting with deionized water for 1 min. 3 ml elution buffer (25 mM Tris,10 mM NaCl, 0.1 % w/v SDS, 100 mM dithiothreitol) was added to each dish and incubated for 1 h at 57 °C. Plates were scraped to harvest the eluate. Harvested proteins were precipitated using 12% (v/v) trichloroacetic acid at 4°C for 30 min. Proteins were pelleted by centrifugation at 5,000g and 4°C for 35 min, followed by another centrifugation round with ice cold acetone for 20 min. The pellets were allowed to dry overnight at 4°C. The protein pellets were then dissolved with buffer containing 6 M urea, 2 M thiourea, 10 mM HEPES (pH 8.0) for in-solution digestion and analyzed *via* mass spectrometry using an LTQ Orbitrap analyser (Thermo Electron). Data was analyzed using the label-free quantification (LFQ) algorithm provided by the MaxQuant Perseus software (version 2.0.6.0).

Statistical analysis

All statistical analysis was characterized by using Prism (GraphPad Software). Unpaired, nonparametric two-tailed Mann-Whitney U-tests were applied to evaluate p-values and statistical significance. p-values lower than 0.05 were considered significant.

Data and code availability statement

The data used in this manuscript are available from the corresponding authors upon reasonable request.

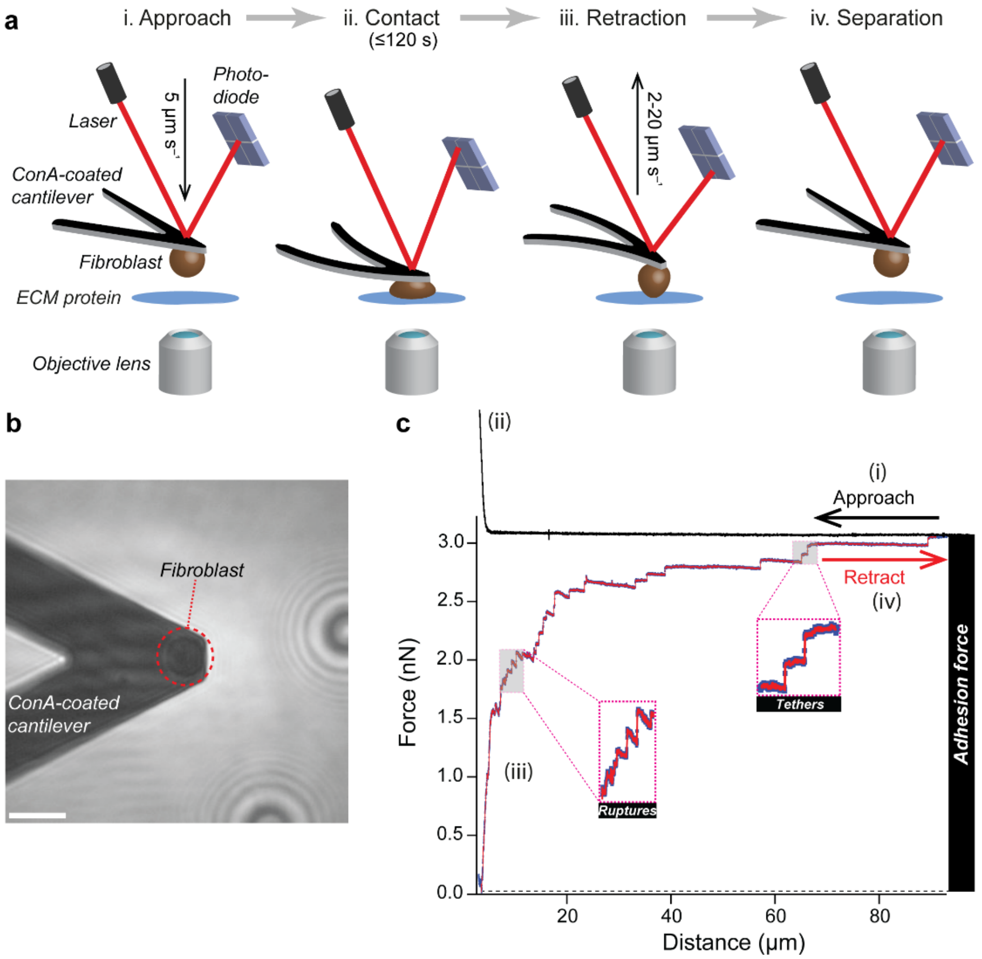


**Extended Data Fig. 1 Experimental scheme used to quantify cell adhesion forces to different substrates under variable retraction speeds. a**, Schematic of atomic force microscopy (AFM)-based single-cell force spectroscopy (SCFS) depicting a single fibroblast attached to a concanavalin A (ConA)-coated cantilever and approached to a vitronectin- or fibronectin-coated glass surface (substrate) (i) until reaching a contact force of 1 nN (ii). After maintaining contact between fibroblast and substrate for a given contact time (here 5, 20, 50, or 120 s) the cantilever was retracted at different retraction speeds in most cases (here 2-20 µm $s^{-1}$) to separate fibroblast and substrate while recording a force-distance curve (iii-iv). **b**, Bright field image of a fibroblast on a ConA-coated cantilever. Scale bar, 20 µm. **c**, A force-distance curve tracing the steps i, ii, iii, and iv depicted in schematic (**a**). The adhesion force is measured by quantifying the maximum cantilever deflection (force) during retraction. Single membrane

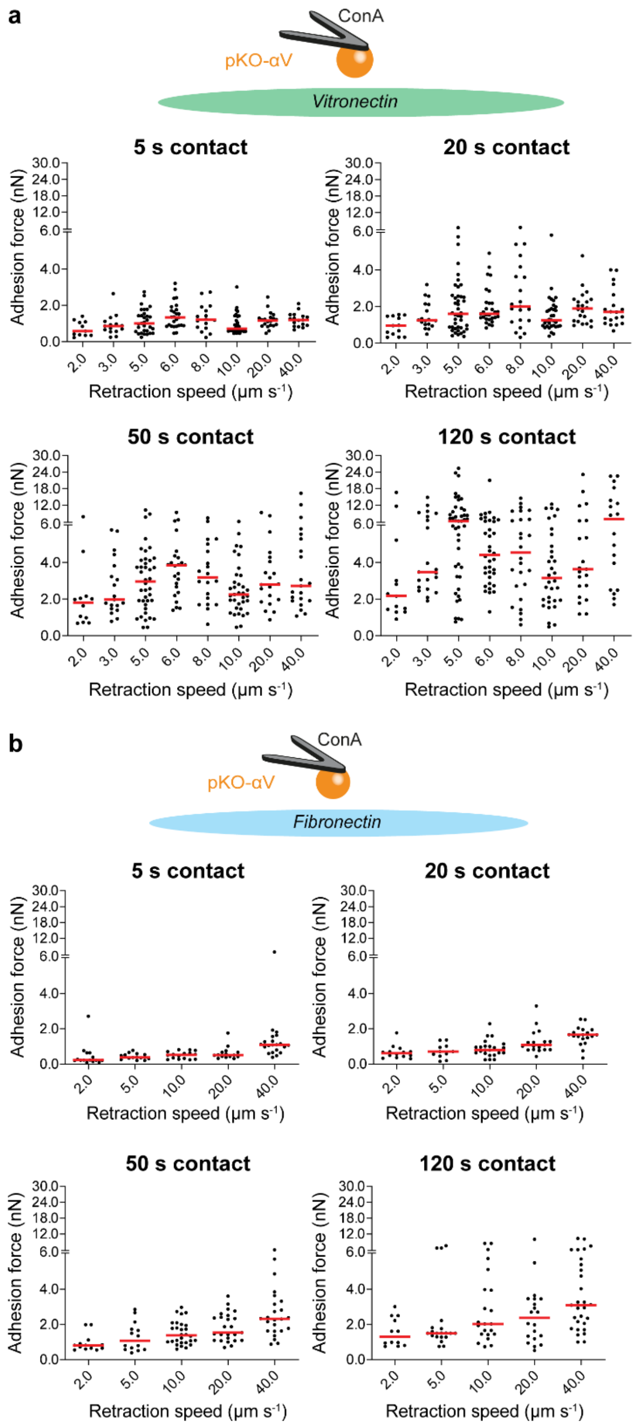


'tether' and integrin-ligand 'rupture' events (inset) are explored in further detail in Extended Data Fig. 5.

**Extended Data Fig. 2 The biphasic adhesion response of fibroblasts is unique to vitronectin and becomes more prominent with increasing contact time. a**, Adhesion force of pKO-αV fibroblasts to vitronectin increases with retraction speed and contact time. The rapid increase in adhesion force, which is observed between 2 μm $s^{-1}$ and 5 μm $s^{-1}$ retraction speed, and the decrease in adhesion force between 5 μm $s^{-1}$ and 10 μm $s^{-1}$ becomes most noticeable at 120 s contact time. **b**, Adhesion force of pKO-αV fibroblasts to fibronectin-coated glass (schematic) increases monophasic with retraction speed and contact time. No rapid increase in adhesion force between 2 μm $s^{-1}$ and 5 μm $s^{-1}$ or a reduction in adhesion force between 5 μm $s^{-1}$ and 10 μm $s^{-1}$ is observed. Dots represent the adhesion force of individual fibroblasts and red bars correspond to medians.

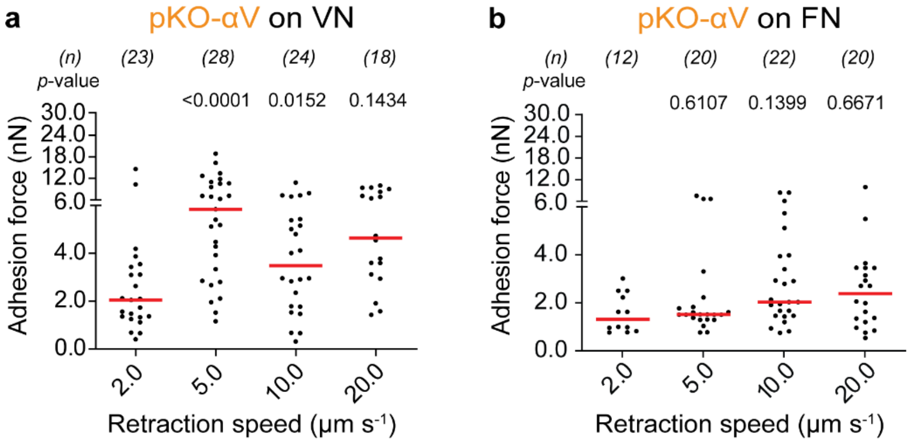


**Extended Data Fig. 3 Adhesion response of pKO-αV fibroblasts at 120 s contact time to vitronectin or fibronectin. a**, pKO-αV fibroblasts adhering to vitronectin (VN) show a biphasic adhesion force in response to retraction speed. This graph has been used as the lightened data in Fig. 2b,d. **b**, Adhesion force of pKO-αV fibroblasts adhering to fibronectin (FN) show a monophasic adhesion force in response to retraction speed. This graph has been used as the lightened data in Fig. 1b and Fig 2d,e. Dots represent the adhesion force of individual fibroblasts and red bars correspond to medians. *p*-values were obtained using two-tailed Mann-Whitney U-test and compare a given retraction speed to the preceding retraction speed (e.g., 5 µm $s^{-1}$ with 2 µm $s^{-1}$). *n* gives the number of fibroblasts measured.

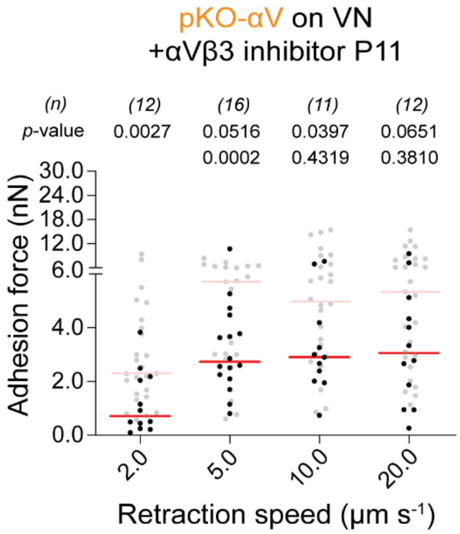


**Extended Data Fig. 4 Chemical inhibition of αVβ3 integrin through P11 verifies that the biphasic adhesion response of pKO-αV fibroblasts to vitronectin (VN) is triggered through αVβ5 integrin.** To verify the αVβ5 integrin specific biphasic cell adhesion response that plateaus at retraction speeds ≥ 5 µm $s^{-1}$ (Fig. 2b), we chemically inhibited αVβ3 integrin using P11 for 30 min. As consequence of this inhibition, we observed a biphasic adhesion response trend similar to that mediated through αVβ5 integrin alone (Fig. 2c), although at lower cell adhesion forces. The data was recorded at 120 s contact time on VN substrate. Dots represent the adhesion force of individual fibroblasts and red bars correspond to medians. *p*-values were obtained using two-tailed Mann-Whitney U-test. *p*-values in the top row compare the data to the pKO-αVβ5 fibroblasts on vitronectin control (lightened) data. *p*-values in the bottom row compare data for a given retraction speed to the preceding retraction speed (e.g., 5 µm $s^{-1}$ with respect to 2 µm $s^{-1}$). *n* gives the number of fibroblasts measured.

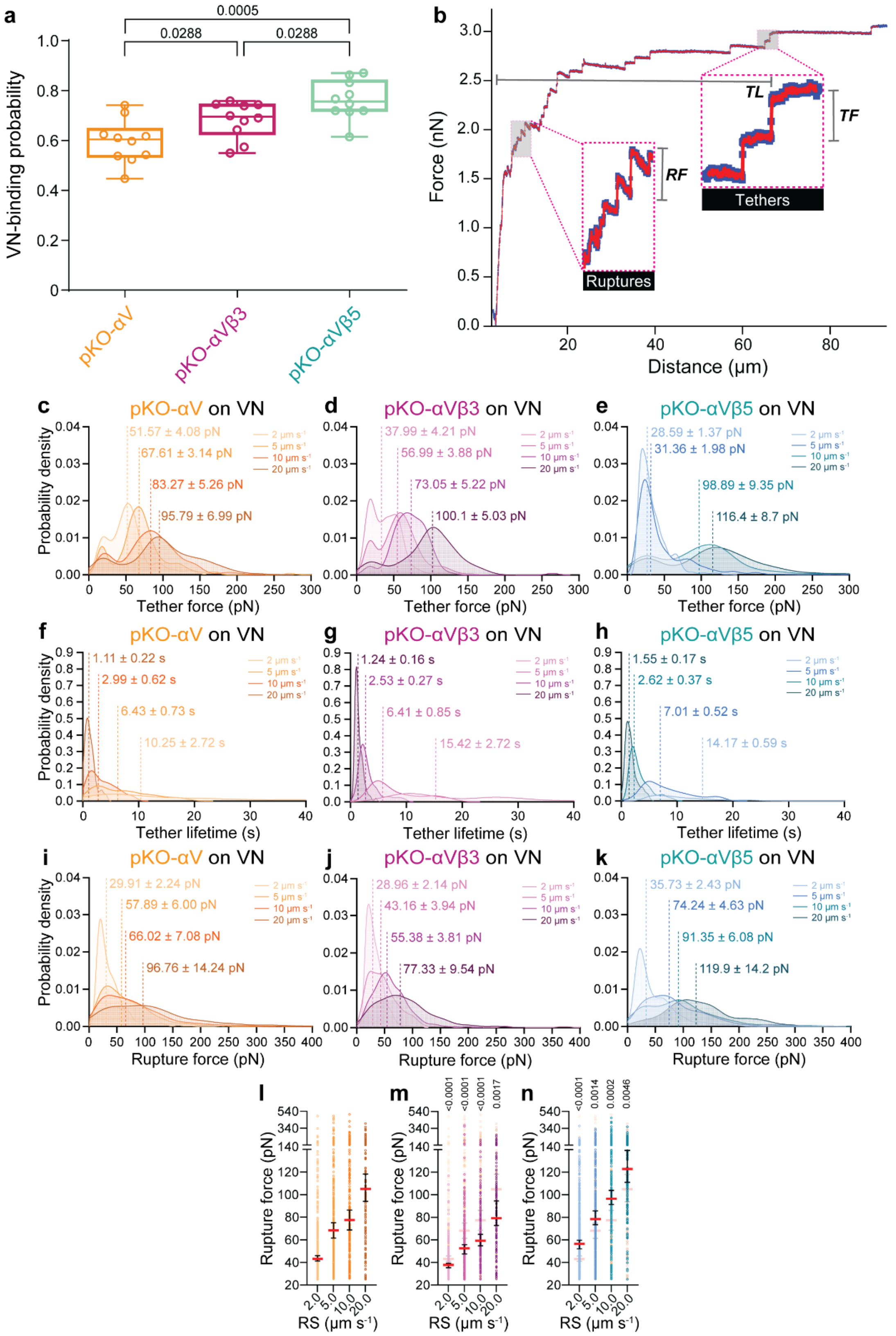


**Extended Data Fig. 5 Ligand-receptor bond characteristics of vitronectin (VN)-bound αV-class integrins influence cellular response to mechanical load. a**, Vitronectin-binding probability of pKO-αV, pKO-αVβ3 and pKO-αVβ5 fibroblasts as measured by repeatedly approaching and retracting a single fibroblast to a vitronectin-coated substrate at 1 $\mu m\ s^{-1}$ velocity and ~50 ms contact time. Each data point represents an outcome from ~100 approach-retraction cycles per cell collected for 10 different fibroblasts. Box plots represent median and interquartile range. *p*-values were obtained using Mann-Whitney U-test. **b**, Schematic of analyzing membrane 'tether' and 'rupture' events (inset) of a force-distance curve recorded upon retracting a fibroblast from vitronectin by SCFS. The membrane 'tether' and 'rupture' events are used to estimate the lifetime and the rupture force of ligand-integrin bonds, respectively [6-8]. *RF* represents rupture force, *TF* tether force, and *TL* tether length. Calculation of tether lifetime is given in Materials and Methods. **c**-**k**, Probability density distribution curves comparing data of 2, 5, 10, and 20 $\mu m\ s^{-1}$ (light to dark colour hue) retraction speeds at 120 s contact time, for tether force (**c**-**e**), tether lifetime (**f**-**h**) and rupture force (**i**-**k**). The dotted lines represent median, and the values denote median± 95% confidence interval (CI) for each speed and condition. The number of 'tether' events analyzed in (**c**) and (**f**) per retraction speed are 128 for 2 $\mu m\ s^{-1}$, 316 for 5 $\mu m\ s^{-1}$, 215 for 10 $\mu m\ s^{-1}$, and 131 for 20 $\mu m\ s^{-1}$, in (**d**) and (**g**) they are 206 for 2 $\mu m\ s^{-1}$, 205 for 5 $\mu m\ s^{-1}$, 106 for 10 $\mu m\ s^{-1}$, and 79 for 20 $\mu m\ s^{-1}$, and in (**e**) and (**h**) they are 830 for 2 $\mu m\ s^{-1}$, 578 for 5 $\mu m\ s^{-1}$, 155 for 10 $\mu m\ s^{-1}$, and 169 for 20 $\mu m\ s^{-1}$. The number of 'rupture' events analyzed per retraction speed in (**i**, **l**) are 575 for 2 $\mu m\ s^{-1}$, 275 for 5 $\mu m\ s^{-1}$, 279 for 10 $\mu m\ s^{-1}$, and 146 for 20 $\mu m\ s^{-1}$, in (**j**, **n**) they are 277 for 2 $\mu m\ s^{-1}$, 240 for 5 $\mu m\ s^{-1}$, 184 for 10 $\mu m\ s^{-1}$, and 136 for 20 $\mu m\ s^{-1}$, and in (**k**, **m**) they are 881 for 2 $\mu m\ s^{-1}$, 371 for 5 $\mu m\ s^{-1}$, 255 for 10 $\mu m\ s^{-1}$, and 136 for 20 $\mu m\ s^{-1}$. Single tether and rupture events were analyzed from SCFS data recorded for Fig. 1b and Fig. 2b,d. **l**, The rupture force of the αV-class integrin-vitronectin bond depends non-linearly on the retraction speed (RS), and thus shows catch bond like behavior in pKO-αV fibroblasts. Data taken from (**i**). **m**, The rupture force of the αVβ3 integrin-vitronectin bonds depends non-linearly on the retraction speed (RS) and shows catch bond like behavior in pKO-αVβ3 fibroblasts. Data taken from (**j**). **n**, The rupture force of the αVβ5 integrin-vitronectin bond bonds depends linearly on the retraction speed (RS) and shows ideal bond like behavior in pKO-αVβ5 fibroblasts. Data taken from (**k**). Red bars in (**l**), (**m**), and (**n**) represent median rupture force with 95% CI. *p*-values in (**m**) and (**n**) were obtained by two-tailed Mann-Whitney U-test comparing data for pKO-αVβ3 or pKO-αVβ5 fibroblasts to pKO-αV fibroblasts, respectively.

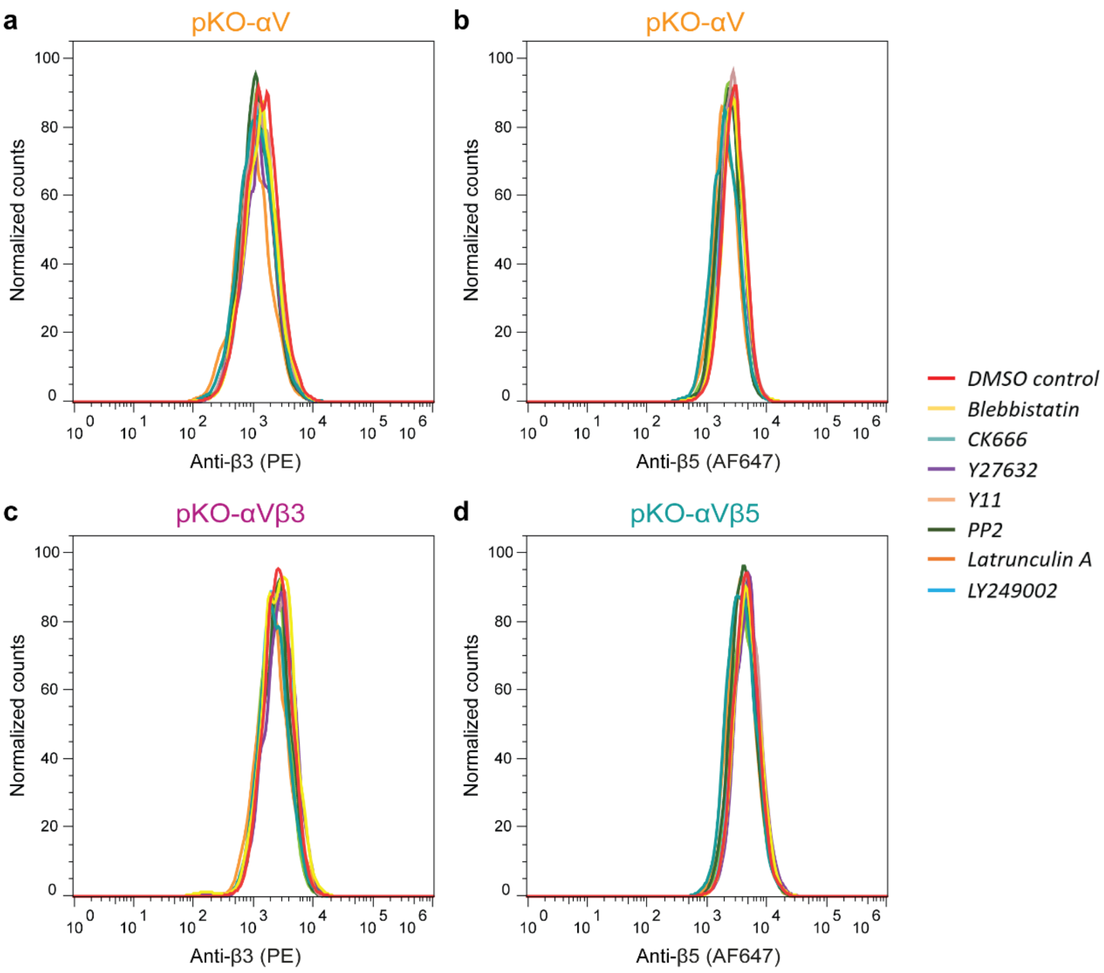


**Extended Data Fig. 6 Integrin β-subunit cell surface expression is independent of small molecule inhibitor treatment.** To confirm the surface expression of integrin β3 and β5-subunit(s) in (**a**, **b**) pKO-αV fibroblasts, (**c**) pKO-αVβ3 fibroblasts, and (**d**) pKO-αVβ5 fibroblasts, the fibroblasts were treated with DMSO (0.1% v/v), 20 µM blebbistatin, 200 µM CK666, 10 µM Y27632, 10 µM Y11, 20 µM PP2, 1 µM latrunculin A, or 10 µM LY249002 for 30 min before being doubly stained with anti-β3 subunit and anti-β5 subunit fluorophore-tagged monoclonal antibodies [control for (Fig. 3) and (Fig. 4)]. **a**, Surface expression of αVβ3 integrin of pKO-αV fibroblasts remains the same for all treatments. **b**, Surface expression of αVβ5 integrin of pKO-αV fibroblasts remains the same for all treatments. **c**, Surface expression of αVβ3 integrin in pKO-αVβ3 fibroblasts remains the same for all treatments. **d**, Surface expression of αVβ5 integrin in pKO-αVβ5 fibroblasts remains the same for all treatments. 5000 fibroblasts were analysed per sample and filtered for live and single cells. The data are representative of two independently repeated experiments.

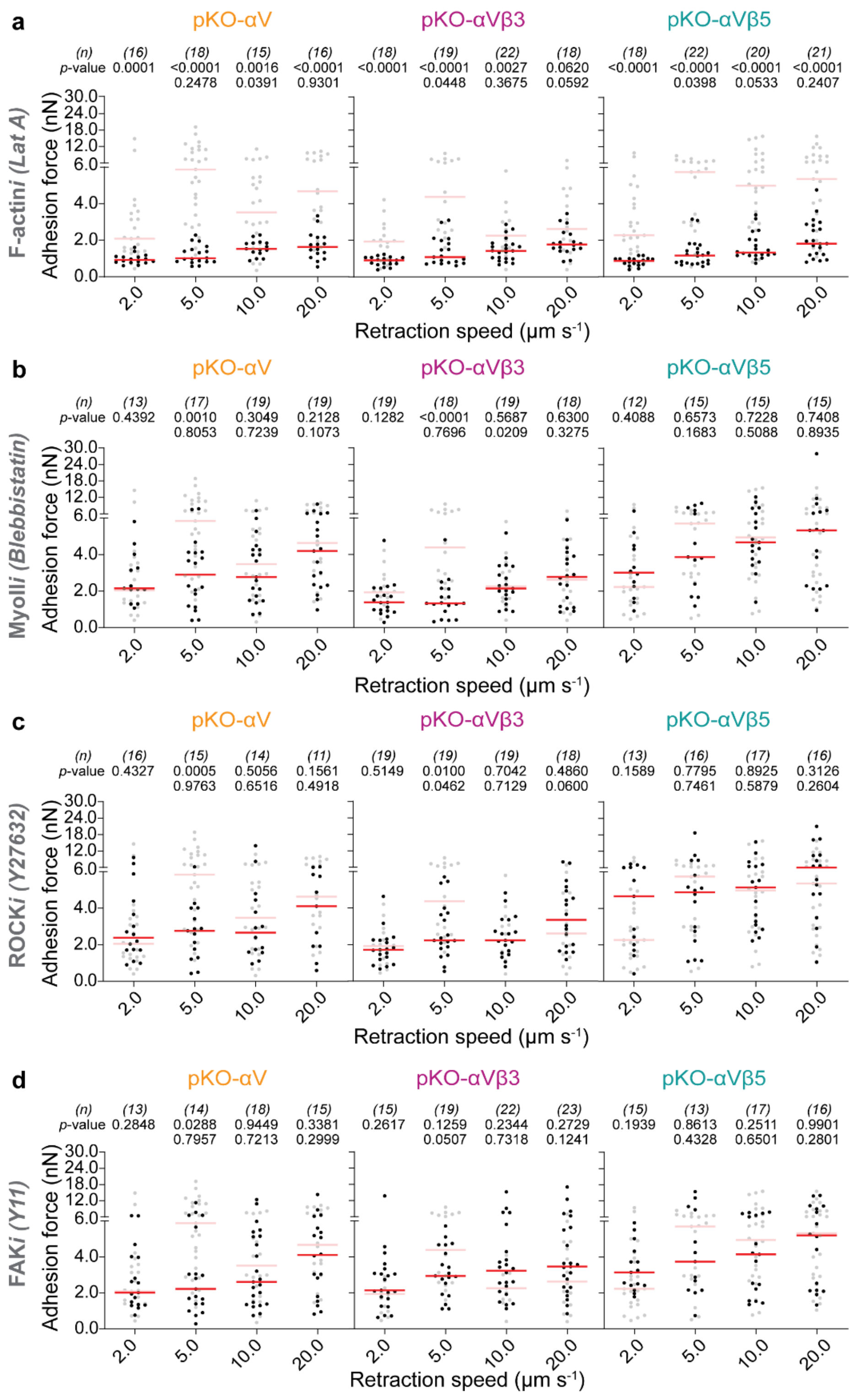


**Extended Data Fig. 7 Integrity and regulation of actomyosin cortex mediates the biphasic response of vitronectin-bound αV-class integrins together with FAK. a-d**, Retraction speed-dependent adhesion force of chemically perturbed pKO-αV, pKO-αVβ3 and pKO-αVβ5 fibroblasts on vitronectin at 120 s contact time. Fibroblasts were treated with 1 μM filamentous actin (F-actin) inhibitor latrunculin A (LatA) (**a**), 20 μM myosin II (MyoII) inhibitor blebbistatin (**b**), 10 µM Rho kinase (ROCK1) inhibitor Y27632 (**c**), and 10 µM focal adhesion kinase (FAK) inhibitor Y11 (**d**). Each dot represents one fibroblast, red bars the median. *p*-values were obtained using two-tailed Mann-Whitney U-test. *p*-values in the top row compare data to unperturbed (lightened) data. *p*-values in the bottom row compare data for a given retraction speed to the preceding speed (e.g., 5 µm $s^{-1}$ with 2 µm $s^{-1}$). *n* denotes the number of fibroblasts measured.

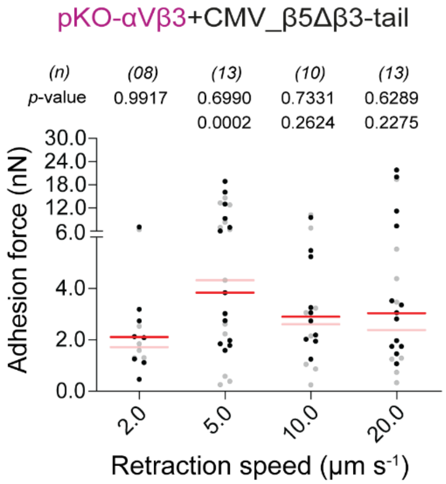


**Extended Data Fig. 8 The tail of the integrin β5-subunit influences early adhesion to vitronectin under mechanical load.** The rescue (as per Extended Data Fig. 4) of the integrin β5-subunit dependent higher cell adhesion force fails if a chimeric β5-subunit[9] (extracellular and transmembrane domain of integrin β5-subunit fused to the cytoplasmic tail of the integrin β3-subunit) is introduced in pKO-αVβ3 fibroblasts. The data was recorded at 120 s contact time on vitronectin substrate. The lightened data represents no rescue of adhesion force occurs if pKO-αVβ3 fibroblasts are transfected with an empty vector (control). The light gray data and red bar show adhesion forces of non-rescued pKO-αVβ3 fibroblasts. Each dot represents one fibroblast, red bars the median. *p*-values were obtained using two-tailed Mann-Whitney U-test. *p*-values in the top row compare the data to the control (lightened) data. *p*-values in the bottom row compare data for a given retraction speed to the preceding retraction speed (e.g., 5 µm $s^{-1}$ with respect to 2 µm $s^{-1}$). *n* gives the number of fibroblasts measured.

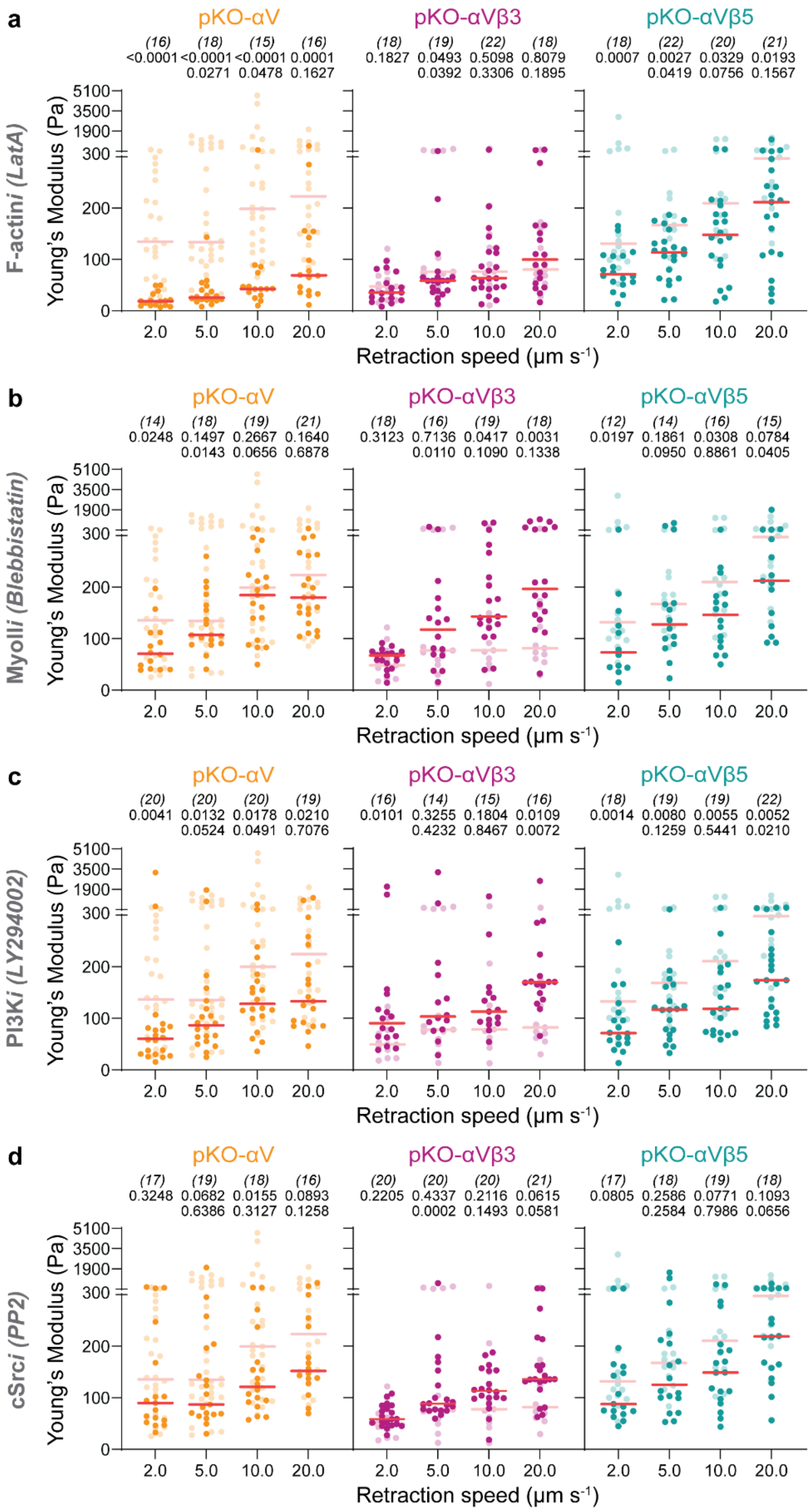


**Extended Data Fig. 9** Integrin β-subunit, Arp2/3 and clathrin mediated endocytosis are key determinants of cytoskeletal stiffening under mechanical load. **a-d**, Retraction speed-dependent apparent cell stiffness (Young's modulus) determined for chemically perturbed pKO-αV, pKOαVβ3 and pKO-αVβ5 fibroblasts on vitronectin substrate after 120 s of contact. Fibroblasts were treated with 1 μM of F-actin inhibitor LatA (**a**), 20 μM myosin II (MyoII) inhibitor blebbistatin (**b**), 10 μM phosphatidyl inositol 3-kinase (PI3K) inhibitor LY294002 (**c**), or 20 μM cSrc kinase inhibitor PP2 (**d**). Each dot corresponds to one retraction force-distance curve corresponding to SCFS data reported in Extended Data Fig.7a,b, Fig. 2b,c, and Fig. 3a,d. Red bars represent median values. *p*-values were obtained using two-tailed Mann-Whitney U-test. *n* denotes the number of cells measured. *p*-values in the top row compare the data to the unperturbed (lightened) data. *p*-values in the bottom row compare data for a given retraction speed to the preceding speed (e.g., 5 μm $s^{-1}$ with respect to 2 μm $s^{-1}$).

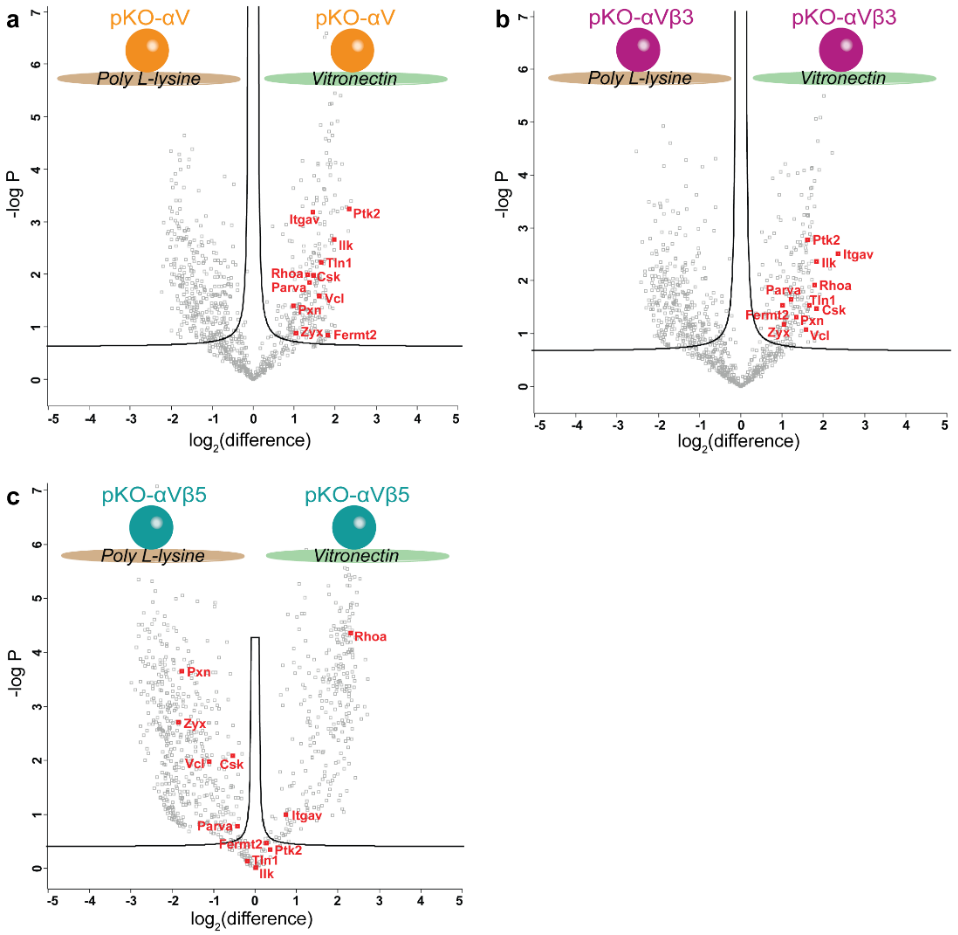


**Extended Data Fig. 10 The consensus adhesome composition of different fibroblasts on vitronectin. a-c**, Consensus adhesome components highlighted in addition to those displayed in Fig. 5, at adhesion sites isolated from pKO-αV fibroblasts adhering to vitronectin or poly-L-lysine (control) for 45 min. Mass spectrometry analysis shows the change in protein abundance on vitronectin (+x axis) against poly-L-lysine (-x axis) for (**a**) pKO-αV fibroblasts, (**b**) pKO-αVβ3 fibroblasts, and (**c**) pKO-αVβ5 fibroblasts. The labelled adhesome components in (**a**) pKO-αV fibroblasts are conserved in (**b**) pKO-αVβ3 fibroblasts, though remarkably altered in (**c**) pKO-αVβ5 fibroblasts. Black lines show the boundaries used to identify protein candidates corresponding to a false-discovery rate of 0.1 and a penalty factor for sample-wide standard deviation of 0.1. The data is representative of four independent replicates.

### References


1. Ran, F.A. *et al.* Genome engineering using the CRISPR-Cas9 system. *Nature Protocols* **8**, 2281-2308 (2013).
2. Montague, T.G., Cruz, J.M., Gagnon, J.A., Church, G.M. & Valen, E. CHOPCHOP: a CRISPR/Cas9 and TALEN web tool for genome editing. *Nucleic Acids Research* **42**, W401-W407 (2014).
3. Friedrichs, J., Helenius, J. & Muller, D.J. Quantifying cellular adhesion to extracellular matrix components by single-cell force spectroscopy. *Nature Protocols* **5**, 1353-1361 (2010).
4. Strohmeyer, N., Bharadwaj, M., Costell, M., Fässler, R. & Müller, D.J. Fibronectin-bound $\alpha5\beta1$ integrins sense load and signal to reinforce adhesion in less than a second. *Nature Materials* **16**, 1262-1270 (2017).
5. Schubert, R. *et al.* Assay for characterizing the recovery of vertebrate cells for adhesion measurements by single-cell force spectroscopy. *FEBS Letters* **588**, 3639-3648 (2014).
6. Krieg, M. *et al.* Atomic force microscopy-based mechanobiology. *Nature Reviews Physics* **1**, 41-57 (2019).
7. te Riet, J. *et al.* Dynamic coupling of ALCAM to the actin cortex strengthens cell adhesion to CD6. *Journal of Cell Science* **127**, 1595-1606 (2014).
8. Viljoen, A. *et al.* Force spectroscopy of single cells using atomic force microscopy. *Nature Reviews Methods Primers* **1**, 63 (2021).
9. Lock, J.G. *et al.* Clathrin-containing adhesion complexes. *Journal of Cell Biology* **218**, 2086-2095 (2019).
